# Steric Gating Dictates Selective Activation of BIRC6 by UBA6 Over UBA1

**DOI:** 10.64898/2025.12.03.691989

**Authors:** Zebin Tong, Rujing Yuan, Xiangwei Wu, Hongyi Cai, Xu Ziyu, Lei Liu, Huasong Ai

## Abstract

The giant E2–E3 chimera BIRC6 is specifically activated by the ubiquitin-activating enzyme UBA6, but not by UBA1. This specificity allows BIRC6 to ubiquitinate downstream substrates (e.g., caspases) and thereby regulate apoptosis. However, the molecular mechanism that underlies UBA6mediated specific activation of BIRC6 remains elusive. Here, we used a Ub^Dha^ (Ub residue 76 dehydroalanine) probe to achieve one-step, activity-based trapping of the transient transthioesterification intermediate, and resolved the cryo-EM structures of two key complexes: 1) the doubly loaded UBA6–BIRC6^UBC^–Ub^T^–Ub^A^ complex (3.4 Å resolution) and 2) the singly loaded UBA6–BIRC6^UBC^–Ub^T^ complex (3.3 Å resolution). Structural analysis reveals that the specific BIRC6–UBA6 pairing is dictated by steric compatibility between an insertion loop (residues 4649– 4653) in BIRC6’s UBC domain and UBA6’s gate helix. Unlike UBA1, UBA6 avoids steric clash with this insertion loop via two key conformational changes: a ~30° clockwise rotation and a 19 Å displacement of its gate helix. These conformational changes collectively create a compatible cavity to accommodate the insertion loop. Biochemical validation confirms two key findings: 1) truncating BIRC6’s insertion loop rescues UBA1-mediated ubiquitin charging of BIRC6^UBC^; 2) swapping UBA6’s gate helix with that of UBA1 abolishes UBA6’s transthioesterification activity towards BIRC6^UBC^. Our study provides key structural insights into the molecular basis of E1–E2 pairing specificity and advances our mechanistic understanding of apoptotic signalling.

## Introduction

Ubiquitin (Ub) and ubiquitin-like proteins (UBLs) modify target proteins to alter their structures, localizations, and interactions, thereby regulating a broad range of cellular functions including protein homeostasis, cell division, and development^1–3^. The covalent attachment of Ub/UBLs to substrates relies on a sequential enzymatic cascade involving E1 activating enzymes, E2 conjugating enzymes, and E3 ligases^4–6^. During E1 activation, the C-terminus of Ub/UBLs is first adenylated in the presence of ATP, followed by thioester bond formation with the catalytic cysteine of E1. The E1 then recruits an E2 conjugating enzyme and transfers the Ub/UBL moiety to the catalytic cysteine of the E2, thereby generating an E2~Ub/UBL thioester intermediate (where “~” denotes a thioester bond), which is a process known as E1–E2 transthioesterification. Finally, E3 ligases transfer Ub/UBLs from the E2 to either their own catalytic cysteine (HECT family) or directly to the target substrate (RING family), depending on their classification^7–9^.

For most UBL systems, conjugation is predominantly initiated by a single dedicated E1 activating enzyme^9^. The ubiquitin system is a notable exception: in 2007, a second human ubiquitin-activating enzyme, UBA6, was identified alongside the canonical E1 enzyme UBA1^10^. UBA6 shares approximately 40% sequence similarity with UBA1^11^. These two E1s activate over 30 different E2 Ub conjugating enzymes, yet they exhibit distinct E2 selectivity^8^. For example, UBA1 specifically activates E2s such as UBE2H, UBE2R2, and UBE2K, whereas UBA6 is specific to E2s including UBE2Z and BIRC6. Additionally, a subset of E2s, such as UBE2G2 and UBE2S, can be activated by both UBA1 and UBA6^10^. This differential E2 preference allows the two E1s to direct ubiquitin to distinct subsets of E3 ligases, thereby mediating the modification of specific substrates and triggering diverse downstream cellular function, including protein degradation, DNA repair, and immune signaling^12^. Previous structural studies of UBA1 in complex with various E2s have delineated the mechanisms underlying ubiquitin adenylation, UBA1~Ub thioester bond formation, and UBA1–E2 transthioesterification^13–21^. In contrast, the molecular basis governing UBA6’s selective recognition of E2 partners has remained unclear, limiting our understanding of E1–E2 pairing specificity.

BIRC6 is a giant 4857-amino acid E2–E3 hybrid enzyme and the only essential inhibitor of apoptosis (IAP) in the apoptotic pathway^22, 23^. Its C-terminus contains a ubiquitin-conjugating (UBC) domain, which is a hallmark of the E2 ubiquitin-conjugating enzyme family^8, 24^. The N-terminal baculoviral IAP repeat (BIR) domains of BIRC6 recognize specific amino acid degrons on substrates, endowing BIRC6 with E3 ligase-like function for direct substrate recruitment^25^. In the apoptosis pathway, BIRC6 is specifically activated by UBA6, but not by UBA1^26–28^. This activation enables BIRC6 to ubiquitinate substrates such as caspases-3, -7, -9 and high-temperature requirement protein A2 (HTRA2), thereby inhibiting apoptosis^29–31^.

Recent structural studies have reported the structures of apo BIRC6 and BIRC6 in complex with various client proteins^26–28, 32^. These structures revealed that BIRC6 forms an unusual, massive headto-tail dimer, thereby assembling into a U-shaped complex with a molecular mass of approximately 1.2 MDa. This architecture features a central cavity flanked by key functional domains such as the UBC and BIR domains. The central cavity serves as a critical binding site for both substrates (including caspases-3, -7, -9, and HTRA2) and the inhibitory protein SMAC (second mitochondria-derived activator of caspases). Although these studies have elucidated the role of BIRC6 as an E3 ligase in substrate recognition and recruitment, the structural and mechanistic basis underlying its specificity for UBA6 (when acting as an E2 enzyme) and its lack of reactivity with UBA1 remains unknown.

In this study, we employed an activity-based Ub^Dha^ probe to achieve one-step capture of the transthioesterification intermediate complex formed by UBA6 and the UBC domain of BIRC6 (BIRC6^UBC^). This strategy enabled the visualization of cryo-EM structures of the complex comprising UBA6, BIRC6^UBC^, and adenylated/thioester-linked ubiquitin. Our structures reveal the molecular mechanism underlying the specific activation of BIRC6 by UBA6 but not UBA1, providing crucial mechanistic insights into the structural basis of E1–E2 pairing specificity.

## Results

### The UBC domain of BIRC6 confers specificity for UBA6

To characterize the enzyme activity of BIRC6, we expressed full-length BIRC6 in HEK293F cells and reconstituted its activity through in vitro transthioesterification reactions. Our results showed that BIRC6 exclusively accepts Ub activated by UBA6, but not by UBA1 (**Fig. 1a**). This result is consistent with previous in vitro reconstitution data and the documented functional synergy between UBA6 and BIRC6 in autophagy regulation^26–28, 33^. We further confirmed that the catalytic UBC domain of BIRC6 (residues 4520-4857) exhibited the same UBA6 preference as full-length BIRC6 (**Fig. 1b**, **Fig. 2a**), indicating that UBA6 E1 enzyme selectivity of BIRC6 originates from its UBC domain. Based on this finding, we utilized the UBC domain of BIRC6 for subsequent mechanistic studies.

**Fig. 1.**
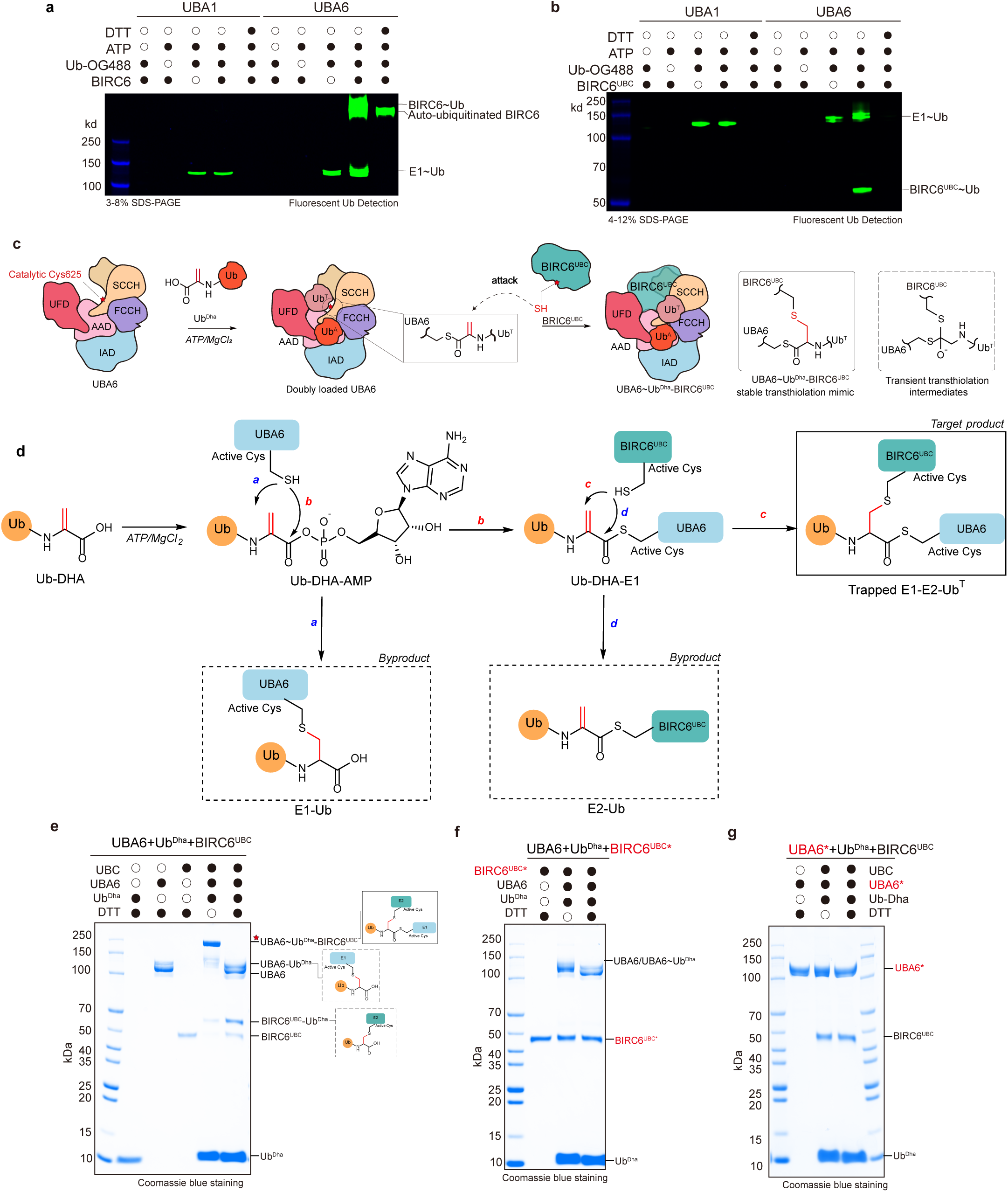
Chemical trapping of the UBA6–BIRC6 transthioesterification reaction. **a-b,** E1-E2 transthioesterification assays for full-length BIRC6 (**a**) and the UBC domain of BIRC6, BIRC6^UBC^ (**b**). The Ub’s transfer was tracked by fluorescently labeled Ub (Ub-OG488). Single asterisk: autoubiquitinated BIRC6. **c**, Schematic illustrating chemical trapping of the UBA6**–**BIRC6^UBC^**–**Ub^T^**–**Ub^A^ complex. Ub^Dha^, a ubiquitin variant with residue G76 replaced by dehydroalanine, can be activated by UbA6 in a manner similar to natural ubiquitin to form doubly loaded UBA6. The electrophilic dehydroalanine then reacts with the active cysteine of the BIRC6^UBC^, generating a trapped transthioesterification mimic. The native and transient transthioesterification intermediate is also shown. **d**, Reaction pathways of Ub^Dha^ in the transthioesterification reaction. Both the target product formation pathway and the byproduct formation pathway are shown. **e-g**, SDS-PAGE analysis of the chemical trapping reaction for wild-type UBA6/BIRC6^UBC^ (**e**), BIRC6^UBC^ with the active-site C4666A mutation (**f**) and UBA6 with the active-site C625A mutation (**g**). The formation of the target product depends on the catalytic cysteines of UBA6 and BIRC6^UBC^.

**Fig. 2.**
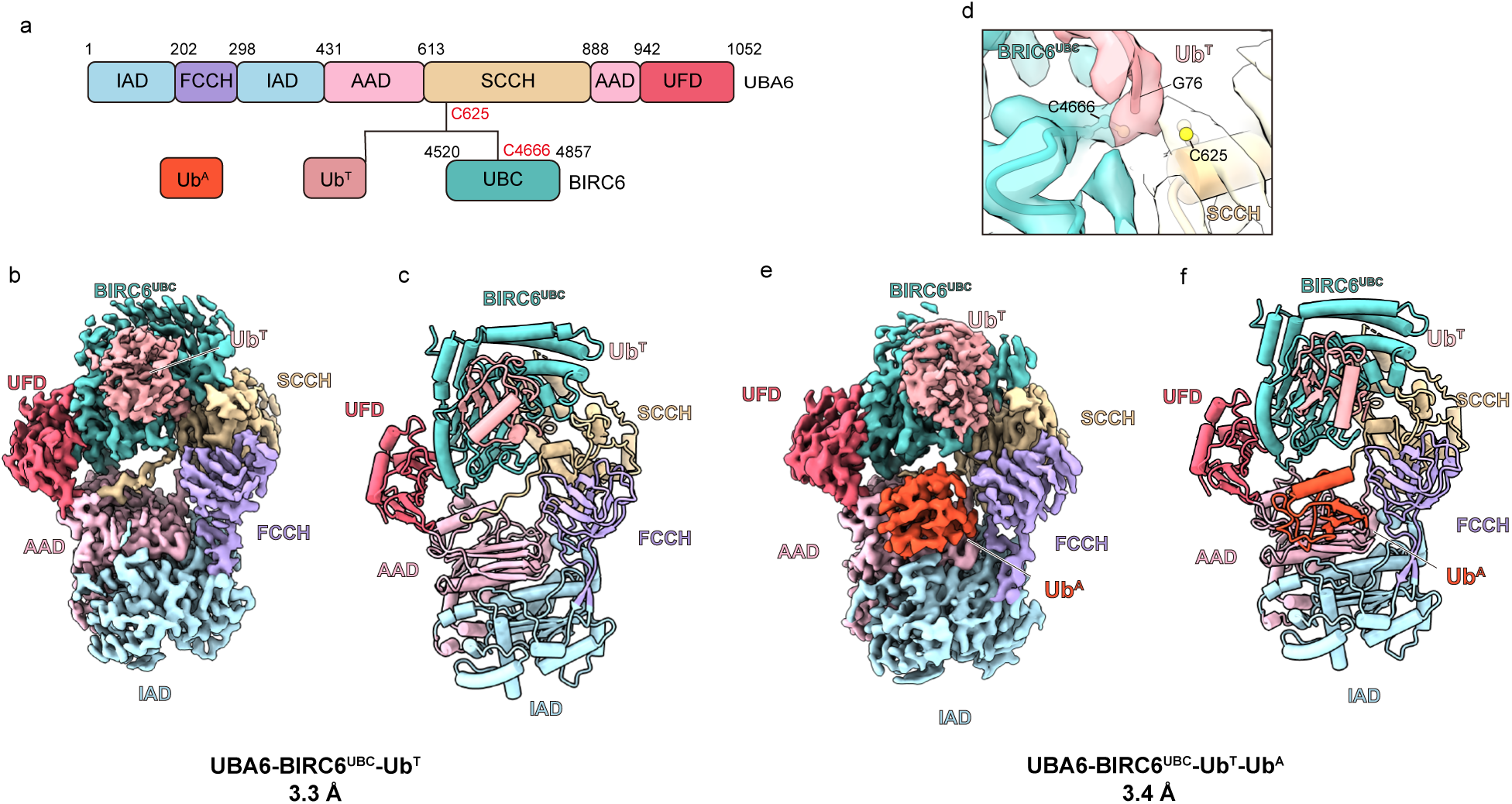
Cryo-EM structures of UBA6–BIRC6 transthioesterification mimic complexes. **a,** Domain diagrams of UBA6, BIRC6^UBC^, Ub^A^, Ub^T^. The domains are colored and labeled accordingly. The black solid line indicates covalent linkages among UBA6, BIRC6^UBC^ and Ub^T^. **b-c,** Cryo-EM map (**b**) and structural model (**c**) of singly loaded UBA6**–**BIRC6^UBC^**–**Ub^T^ complex. Each domain is labeled and the color scheme is the same as in (**a**). **d,** A close-up view shows the local density and the fitted model at the transthioesterification center. **e-f,** Cryo-EM map (**e**) and structural model (**f**) of doubly loaded UBA6**–**BIRC6^UBC^**–**Ub^T^**–**Ub^A^ complex.

### One-step trapping of the E1-E2-Ub^T^-Ub^A^ complex

During the E1-E2 transthioesterification process, E1 catalyzes the adenylation of ubiquitin (Ub^A^), which is then transferred to the catalytic cysteine of E1 to form a thioester-linked ubiquitin (Ub^T^). The E1 subsequently recruits and adenylates a second ubiquitin molecule, resulting in a doubly loaded E1 complex with a 1:2 E1-to-ubiquitin stoichiometry^34, 35^. The doubly loaded E1-E2- Ub^T^- Ub^A^ complex exhibits maximal transfer efficiency of Ub^T^ to E2’s catalytic cysteine, establishing this state as the definitive active conformation for E1-E2 ubiquitination. We sought to determine the structure of the doubly loaded E1-E2-Ub^T^-Ub^A^ complex to enhance our understanding of the molecular mechanisms underlying the specificity of the E1-E2 pair. However, capturing the structure of the doubly loaded E1-E2-Ub^T^-Ub^A^ complex is challenging due to the unstable nature of the thioester bond, the low affinity between E1 and E2, and the transient nature of the transthioesterification intermediates.

The atomic-level precision of protein chemical synthesis^36–41^ has enabled the rational design of chemical trapping strategies, including mechanism-inspired and activity-based probes, to advance the mechanistic study of ubiquitin-related enzymes^42, 43^. These ubiquitin probes have significantly advanced structural studies of ubiquitin-related enzymes, including the ubiquitin E1-E2-E3 cascade^14, 15, 19–21, 42, 44–53^ and deubiquitinating enzymes (DUBs)^54–56^. Here, we employed the activity-based probe Ub^Dha^ probe to chemically capture the doubly loaded E1–E2–Ub^T^–Ub^A^ complex^57^. Ub^Dha^ is commercially available and can be synthesized by recombinantly expressing UbG76C and converting the cysteine residue to the Dha moiety using 2,5-dibromohexanediamide in a one-step chemical reaction (**Extended Fig. 1a-c)**^57^. Although the residue 76 of Ub^Dha^ is replaced by the electrophilic dehydroalanine, it remains recognizable and can be catalyzed by the natural ubiquitin machinery^57^. Ub^Dha^ can be activated at its C-terminus by E1 in a manner similar to natural ubiquitin, leading to the formation of a thioester-linked product E1~Ub^Dha^ intermediate, which can subsequently transfer to E2s (**Fig. 1c-d**). Moreover, the electrophilic dehydroalanine of Ub^Dha^ can engage in an irreversible reaction with the active cysteine residues of E1, E2, HECT/RBR-type E3 ligases and deubiquitinases^54–56^.

We next biochemically tested the utility of the Ub^Dha^ probe for covalent trapping of the UBA6– BIRC6^UBC^ complex. To this end, we mixed the Ub^Dha^ probe, UBA6, BIRC6^UBC^, and ATP/Mg^2+^ while optimizing the concentrations and molar ratios of UBA6, BIRC6^UBC^, and Ub^Dha^. Our results showed that at the optimized molar ratio of UBA6:BIRC6^UBC^: Ub^Dha^ = 1:1:12.5, with UBA6 at a final concentration of 1 μM, nearly all UBA6 was cross-linked with BIRC6^UBC^ and Ub^Dha^ to form the doubly loaded UBA6-BIRC6^UBC^-Ub^T^-Ub^A^ complex (171kD) (**Fig. 1e, lane 4; Extended Fig. 1d**). The crosslinked byproducts UBA6-Ub^Dha^ (126kD) and BIRC6^UBC^~Ub^Dha^ (43kD) can be removed through size-exclusion chromatography, allowing for the isolation of high-purity doubly loaded UBA6-BIRC6^UBC^Ub^T^-Ub^A^ complexes (**Extended Fig. 1e, f)**. Additionally, mutating the active site C4666 of BIRC6^UBC^ or C625 of UBA6 to alanine resulted in negligible cross-linking bands, indicating activity-based specific cross-linking (**Fig. 1f, g**). It is important to note that this Ub^Dha^ method is activity-based sitespecific cross-linking, requiring no point mutations or modifications of E1 and E2.

### Structure of the UBA6-BIRC6^UBC^-Ub transthioesterification mimic

The vitrified samples were analyzed by cryo-EM, yielding two reconstructions of the singly loaded UBA6–BIRC6^UBC^–Ub^T^ complex and the doubly loaded UBA6–BIRC6^UBC^–Ub^T^–Ub^A^ complex at resolutions of 3.3 Å and 3.4 Å, respectively (**Fig. 2b, e, Extended Fig. 2, 3a**). Both constructs exhibited the identical overall architecture, with BIRC6^UBC^ sandwiched between the UFD and SCCH domains of UBA6. The difference between these reconstructions was the presence of visualized density for the Ub^A^. The presence of both complexes can be explained by the differential stability of their ubiquitin moieties. Ub^T^ forms a covalent thioester linkage with UBA6-Cys625, whereas Ub^A^ is bound non-covalently and is thus inherently more dynamic. This lability makes Ub^A^ prone to dissociation during sample processing, including size-exclusion chromatography and cryo-EM vitrification. The Cryo-EM map unambiguously reveals clear density connecting the catalytic residues UBA6 Cys625, BIRC6^UBC^ Cys4666, and Ub^T^ Gly76, validating our trapping strategy (**Fig. 2d**).

Both reconstructions exhibit well-ordered UBA6 and BIRC6^UBC^, with local resolutions ranging between 2.9–3.4 Å. By contrast, Ub^T^ is comparatively disordered, with a local resolution ranging from 4.0 to 5.8 Å (**Extended Fig. 3b**). Focused 3D classification on Ub^T^ for both the singly loaded UBA6–BIRC6^UBC^–Ub^T^ and doubly loaded UBA6–BIRC6^UBC^–Ub^T^–Ub^A^ complexes yielded 10 reconstructions each, revealing pronounced Ub^T^ heterogeneity (**Extended Fig. 3c)**. These reconstructions capture a conformational continuum—from Ub^T^ positioned near UBA6’s FCCH domain to states proximal to BIRC6^UBC^. This spatial progression implies the process of Ub^T^ transfer from UBA6 catalytic cystine to BIRC6^UBC^ (**Extended Fig. 3)**.

Structural models were obtained for the singly loaded UBA6–BIRC6^UBC^–Ub^T^ and doubly loaded UBA6–BIRC6^UBC^–Ub^T^–Ub^A^ complexes via structure docking, manual refinement, and Phenix realspace refinement (**Fig. 2c, f, Extended Fig. 4, 5)**. Given that the doubly loaded UBA6–BIRC6^UBC^– Ub^T^–Ub^A^ complex exhibits maximal transfer efficiency of Ub^T^ to E2’s catalytic cysteine, subsequent analysis focuses on the doubly loaded UBA6–BIRC6^UBC^–Ub^T^–Ub^A^ complex.

### Combinatorial recognition of BIRC6^UBC^ by UBA6

The 40 human E2 enzymes share a conserved catalytic domain of approximately 150 amino acid residues, which is termed the UBC domain. UBC domain comprises four α-helices and a four-stranded β-sheet connected by β-loops (**Extended Fig. 6a)**. Beyond the canonical UBC scaffold, the UBC domain of BIRC6 features N-terminal/C-terminal extensions and internal insertions (such as α3 insertion and insertion loop) that constitute structural elements critical for UBA6-BIRC6^UBC^ recognition beyond the core UBC scaffold (**Extended Fig. 6b-d)**. Therefore, in contrast to other E1E2 interactions, the molecular recognition between BIRC6^UBC^ and UBA6 is combinatorial: BIRC6^UBC^ engages with the UFD, AAD, and SCCH domains of UBA6 through six distinct interfaces. This binding buries approximately 1,400 Å^2^ of the BIRC6^UBC^ surface area within the complex (**Fig. 3e**).

**Fig. 3.**
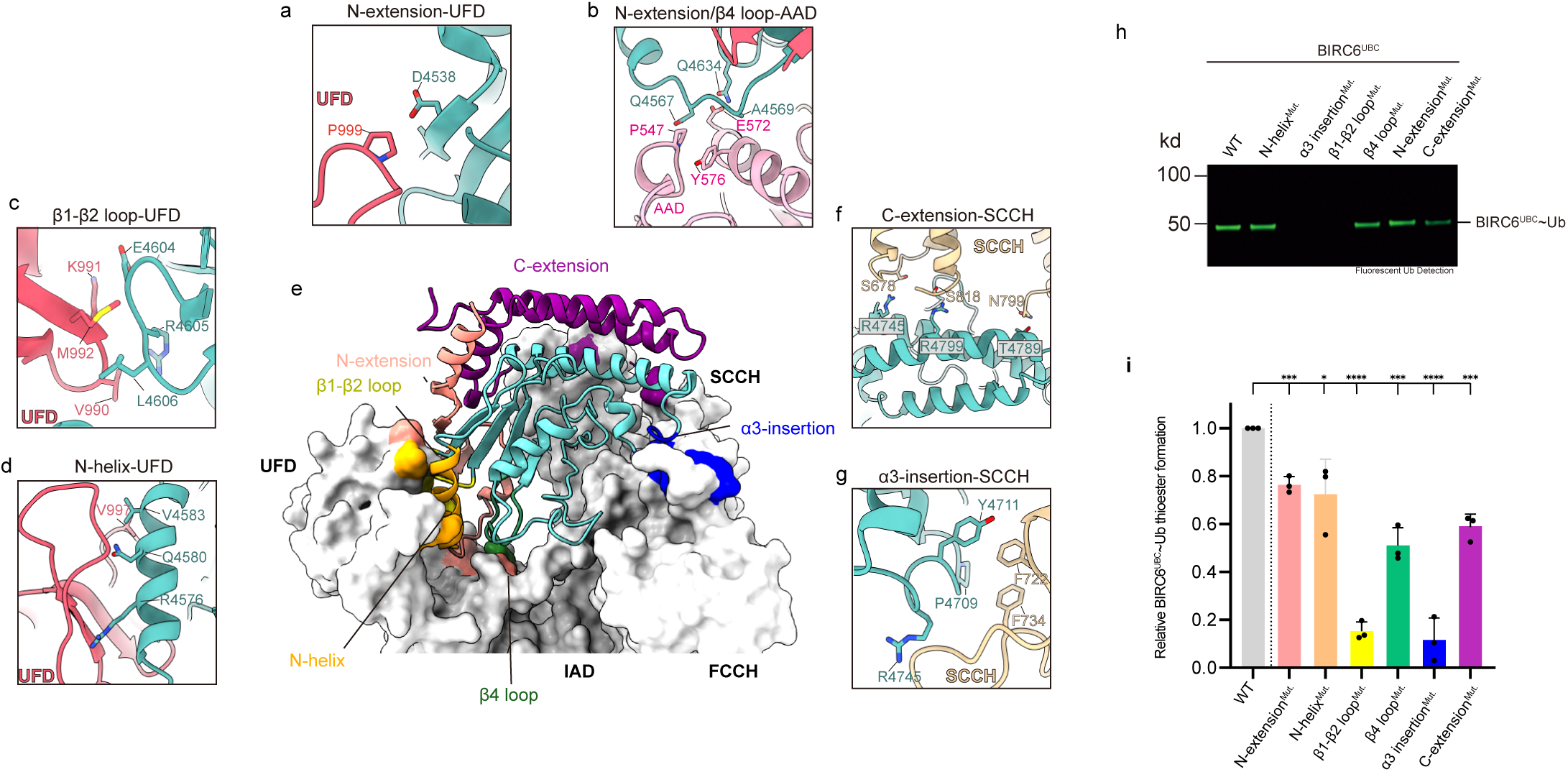
Molecular recognition of BIRC6^UBC^ by UBA6. **a**, Close-up view of the interface between Nextension of BIRC6^UBC^ and the UBA6 UFD domain. The interacting residues are labeled. **b**, Close-up view of the interface between N-extension/β4 loop of BIRC6^UBC^ and the UBA6 AAD domain. **c**, Closeup view of the interface between the β1-β1 loop of BIRC6^UBC^ and the UBA6 UFD domain. **d**, Closeup view of the interface between the N-helix of BIRC6^UBC^ and the UBA6 UFD domain. **e,** Overall view of the contacts between BIRC6^UBC^ and UBA6. UBA6 is shown in surface representation, and BIRC6^UBC^ in cartoon representation. The different regions of BIRC6^UBC^ are colored and labeled. **f**, Close-up view of the interface between the C-terminal extension of BIRC6^UBC^ and the UBA6 SCCH domain. **g**, Close-up view of the interface between α3-insertion of BIRC6^UBC^ and the UBA6 SCCH domain. **h**, In vitro transthioesterification assays using fluorescently labeled Ub. Various BIRC6^UBC^ mutants were analyzed. The gel image represents independent biological replicates (n = 3). **i**, Bar graph showing the fraction of BIRC6^UBC^~Ub presented as mean ± s. d. from n = 3 independent experiments. Each data point is shown as a black dot (data from **h**). A two-sample, two-tailed Student’s unpaired ttest was used to calculate p values; ****p < 0.0001, ***p < 0.001, *p < 0.05.

The first interface involves the N-terminal extension of BIRC6^UBC^ engaging UBA6’s UFD and AAD domains. The N-terminal extension of BIRC6^UBC^, positioned directly above the UBC domain, traverses the cleft between UFD and AAD and winds back to the N-helix of BIRC6^UBC^, forming multiple hydrophobic interactions involving both side chains and backbone of UBA6 (**Fig. 3e**). Specifically, UFD residue P999 is spatially proximate to the backbone of BIRC6^UBC^ residue D4538 (**Fig. 3a, Extended Fig. 7a**), the backbone of BIRC6^UBC^ residue A4569 is near that of AAD residue Y576, and the AAD residue P547 is adjacent to that of backbone of BIRC6^UBC^ residue Q4567. These spatial relationships suggest potential hydrophobic interactions (**Fig. 3b, Extended Fig. 7b**). The second and third interfaces involve interactions between the BIRC6^UBC^ N-helix/β1-β2 loop and UFD (**Fig. 3c, d, Extended Fig.7c, d**). Regarding the BIRC6^UBC^ N-helix/UFD interface, BIRC6^UBC^ residues R4576 and Q4580 are within hydrogen bonding distance of UFD’s backbone carbonyl group. Furthermore, BIRC6^UBC^ residue V4583 may potentially participate in a hydrophobic interaction with UFD residue V997 (**Fig. 3b, Extended Fig. 7d**). In the β1-β2 loop/UFD interface, a potentially van der Waals network connects the backbone frameworks of β1-β2 loop residues E4604- R4605- L4606 and UFD residues V990- K991- M992 (**Fig. 3c, Extended Fig. 7c**). Fourth, the residue Q4634 in the β4 loop of BIRC6^UBC^ may form a salt bridge with residue E572 in the AAD domain (**Fig. 3b, Extended Fig. 7b**). Fifth, the α3 insertion residues P4709 and Y4711 of BIRC6^UBC^ may interact with residues F722 and F734 in the SCCH domain through Van der Waals forces, while residue R4745 of BIRC6^UBC^ is within hydrogen bonding distance of the backbone atoms (**Fig. 3g, Extended Fig. 7e**). The sixth interaction interface involves the C-terminal extension of BIRC6^UBC^ and the SCCH domain. The residues R4745, R4799, and T4789 in the C-terminal extension of BIRC6^UBC^ are within hydrogen bonding distance of the backbone of residues S678 and S818, as well as residue N799 in the SCCH domain (**Fig. 3f, Extended Fig. 7f**).

To validate these interactions, we conducted E1-E2 charging assays using BIRC6^UBC^ with mutations at six different interfaces (**Fig. 3h, i**). The mutants included: an N-terminal extension mutant (A4569R); an N-helix mutant (R4576A-Q4580A-V4583G); a β1-β2 loop mutant (E4604R-R4605GL4606G); a β4 loop mutant (Q4634A); an α3-insertion mutant (P4709R-Y4711G-R4745A); and a Cterminal extension mutant (R4745A-R4799A-T4789A). The results indicate that mutations in the Nterminal extension and N-helix of BIRC6^UBC^ reduce the activity of charged Ub compared to the wild type (WT) by approximately 25%. In contrast, mutations in the β4 loop and C-terminal extension result in a 50% reduction in activity, while mutations in the β1-β2 loop and α3 insertion almost abolish activity.

### Molecular basis of UBA6 specificity for BIRC6^UBC^

To elucidate the molecular mechanisms underlying the UBA6 specificity of BIRC6^UBC^, we compared our resolved structures of UBA6–BIRC6^UBC^–Ub^T^–Ub^A^ complexes with the previously determined UBA1–E2–Ub structures (PDB: 9B5E, 9B5N, 4II2, 5KNL, 7K5J) (**Extended Fig. 8a**) ^13, 14, 16, 17^. The alignment revealed highly similar overall conformations with RMSDs of 1.3-1.7 Å among these structures (**Extended Fig. 8a**). Further comparison of the doubly loaded UBA6–BIRC6^UBC^– Ub^T^–Ub^A^ structure with the recently determined SpUBA1–Ubc4–Ub^T^–Ub^A^ complex (PDB: 9B5E) identified a critical structural distinction: although BIRC6^UBC^ and Ubc4 occupy equivalent positions on their respective E1 enzymes, a pronounced steric clash occurs between BIRC6^UBC^ and SpUBA1 (**Fig. 4a, b**)^13^. This spatial clash was consistently observed in other aligned UBA1–E2–Ub structures, involving the BIRC6^UBC^ amino acids 4649-4653 (referred to as the insertion loop) and the α22-α23 helix of spUBA1 (referred to as the UBA1 Gate helix, amino acids 632-656) (**Fig. 4c-e**). Sequence alignment indicates that BIRC6^UBC^ has a five-amino-acid insertion loop compared to spUbc4 (and its human homolog UbcH5a), which creates a spatial conflict with the gate helix of spUBA1 (**Fig. 4d, g**). In contrast, the gate helix of UBA6 (amino acids 666-690) avoids this steric clash through a 30° clockwise rotation and 19 Å movement of its α22-helix, accommodating the insertion loop of BIRC6^UBC^ (**Fig. 4f**).

**Fig. 4.**
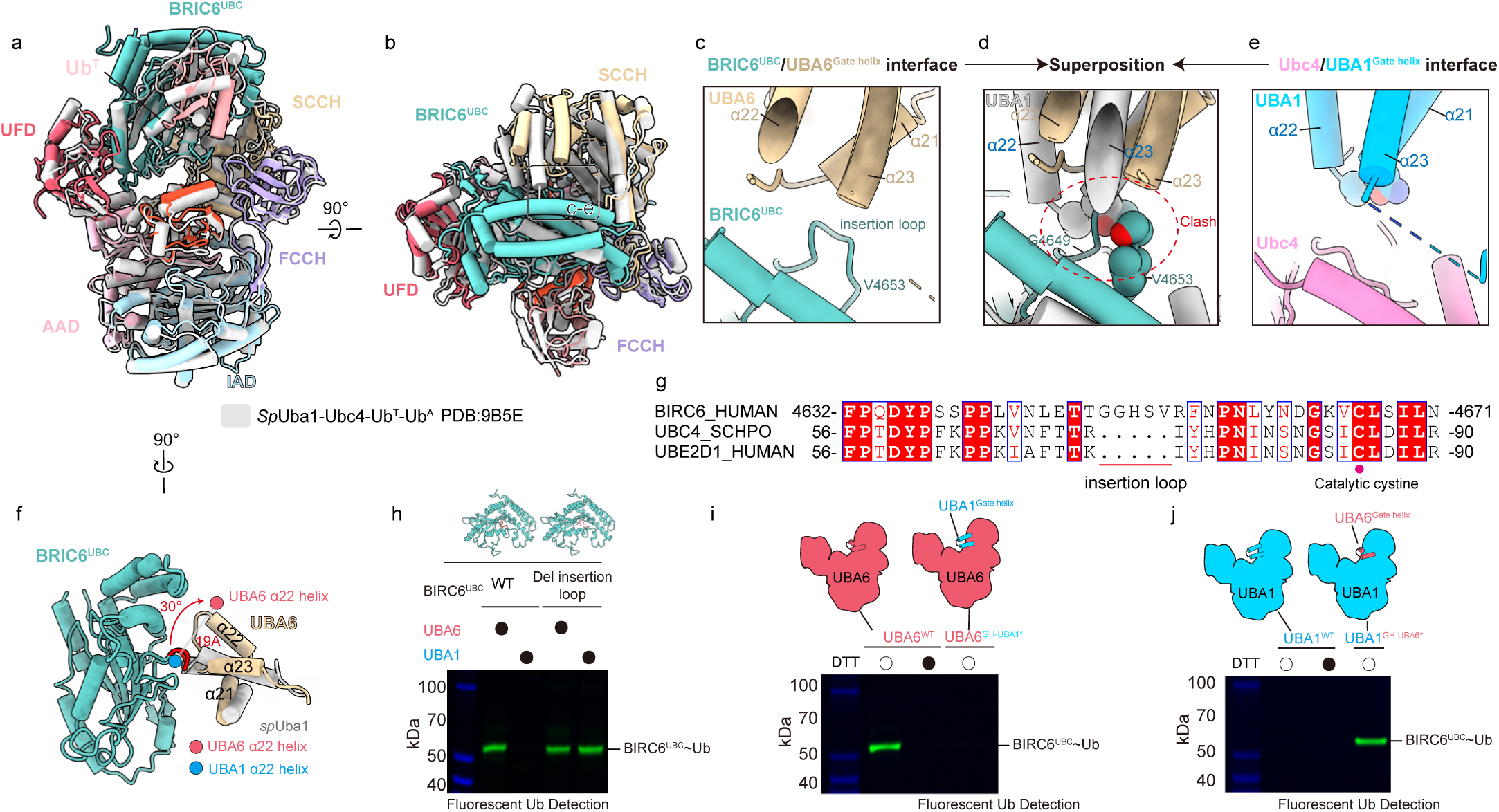
Structural identification and biochemical validation of the gate helix and insertion loop that determines UBA6–BIRC6^UBC^ pairing specificity. **a-b,** Comparison of the doubly loaded UBA6**–**BIRC6^UBC^**–**Ub^T^**–**Ub^A^ complex with representative doubly loaded spUBA1–Ubc4-Ub^T^**–**Ub^A^ complex structures (9B5E) revealed a pronounced steric clash between BIRC6^UBC^ and spUBA1. The rounded rectangle region corresponds to the close-up view shown in **c-e. c**, Close-up view of the spatial arrangement between BIRC6^UBC^ and the UBA6 gate helix (α22-α23 helix). The BIRC6^UBC^ (4649-4653) is shown and labeled as the insertion loop. **d**, Structural alignment revealed a steric clash between the spUBA1 gate helix (α22–α23 helix, amino acids 632-656) and the insertion loop of BIRC6^UBC^. **e**, Close-up view of the spatial arrangement between Ubc4 and the spUBA1 Gate helix (α22-α23 helix, amino acids 632-656). **f**, Relative to the α22 helix (amino acids 632-641) in UBA1, the α22 helix (amino acids 666-674) in UBA6 is rotated 30° clockwise and shifted by 19 Å, generating a space that accommodates the insertion loop of BIRC6^UBC^. **g,** Sequence alignment shows BIRC6 contains a five– amino-acid insertion loop compared with other representative E2 enzymes (spUbc4 and hsUbE2D1). **h**, Truncation of the BIRC6^UBC^ insertion loop restores UBA1-mediated Ub charging to a level comparable to that of UBA6. **i-j**, Chimeric E1 enzymes were generated by swapping the Gate helix between UBA1 and UBA6. UBA6^GH-UBA1*^, in which the gate helix of UBA6 was replaced with that of UBA1, loses transthioesterification activity toward BIRC6^UBC^ (**i**), whereas UBA1^GH–UBA6*^, carrying the gate helix of UBA6, restores Ub charging activity with BIRC6^UBC^. The gel image represents independent biological replicates (n = 3).

To validate whether steric clashes prevent UBA1-activated ubiquitin transfer to BIRC6^UBC^, we truncated the insertion loop of BIRC6^UBC^, then measured its transthioesterification activity towards UBA1 and UBA6. Deletion of the insertion loop of BIRC6^UBC^ rescued UBA1-mediated ubiquitin charging, restoring activity comparable to UBA6 levels (**Fig. 4h**). Next, we engineered chimeric E1 enzymes through gate helix exchange. UBA6 harboring UBA1’s gate helix (referred to as UBA6^GHUBA1*^) retained ubiquitin activation activity comparable to wild-type UBA6 but completely lost transthioesterification activity with BIRC6^UBC^ (**Fig. 4i**), whereas UBA1 bearing UBA6’s gate helix (UBA1^GH-UBA6*^) rescued ubiquitin charging activity with BIRC6^UBC^ (**Fig. 4i, j**).

Collectively, gate helix divergence governs E1-E2 pairing specificity between UBA1 and UBA6. A 30° rotation in the UBA6 gate helix avoids steric clash with the BIRC6^UBC^ insertion loop, while the UBA1 gate helix imposes spatial conflict, thus establishing exclusive BIRC6^UBC^ selectivity for UBA6.

### Gating mechanism for E1-E2 Pair specificity in transthioesterification

Previous studies established a model for the E1-E2 transthioesterification cycle: A doubly loaded E1–Ub^T^-Ub^A^ recruits an E2 to its UFD domain in a distal configuration. The UFD then rotates toward the E1 active site, positioning the E2 catalytic cysteine for Ub^T^ transfer and subsequent E2~Ub thioester bond formation^14, 17, 18^. To determine whether the steric clash with the gate helix impairs BIRC6^UBC^ binding to UBA1 or specifically blocks the subsequent transthioesterification reaction, we performed microscale thermophoresis (MST) binding assays to compare the binding affinity of BIRC6^UBC^ for UBA1 versus UBA6. The result suggested that BIRC6^UBC^ binds UBA1 and UBA6 with comparable affinities (13 μM and 23 μM, respectively) (**Fig. 5a**). Furthermore, titrating UBA1 into transthioesterification reactions containing BIRC6^UBC^ and UBA6 inhibited formation of BIRC6^UBC^~Ub (**Fig. 5b, c**). These results confirm that although UBA1 binds BIRC6^UBC^, it cannot catalyze ubiquitin transfer, ruling out the hypothesis that the gate helix prevents UBA1 from binding BIRC6^UBC^.

**Fig. 5.**
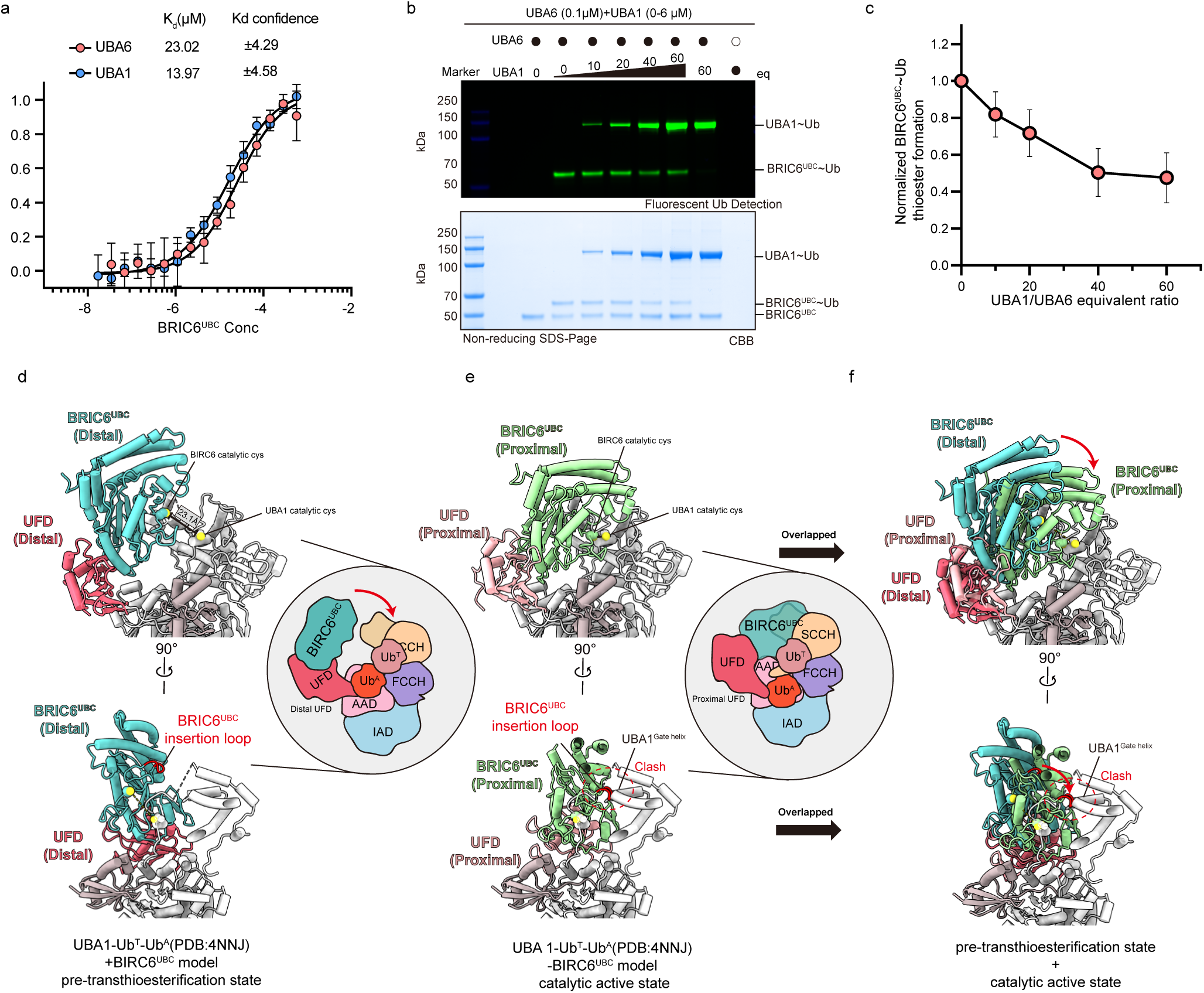
The steric block mediated by the UBA1 gate helix/BIRC6^UBC^ insertion loop prevents transthioesterification despite comparable UBA1–BIRC6^UBC^ binding. **a**, Microscale thermophoresis (MST) binding assays showed comparable affinities between BIRC6^UBC^ and UBA1 or UBA6 (K_d_ = 13 μM and 23 μM, respectively). **b-c**, Increasing concentrations of UBA1 were titrated into transthioesterification reactions containing UBA6 and BIRC6^UBC^, with Mg^2+^, ATP, and fluorescently labeled Ub present in excess. The formation of BIRC6^UBC^~Ub was progressively inhibited, as shown by SDS–PAGE (**b**) and quantified in (**c**) as the mean ± s.d. of three independent experiments. **d–f**, Structural modeling suggests that the gate helix of UBA1 sterically blocks BIRC6^UBC^ from adopting the catalytic transthioesterification conformation. **d,** BIRC6^UBC^ (from the UBA6–BIRC6^UBC^–Ub^T^–Ub^A^ structure) was docked into the doubly loaded UBA1–Ub^T^–Ub^A^ complex (PDB: 4NNJ) by aligning their UFD domains, generating a pre-transthioesterification model in which the active Cys of UBA1 and BIRC6^UBC^ are >23 Å apart. **e,** BIRC6^UBC^ (from the UBA6–BIRC6^UBC^– Ub^T^–Ub^A^ structure) was docked into the doubly loaded spUBA1-Ubc4–Ub^T^–Ub^A^ complex (PDB: 9B5E) by aligning E2 to generate the catalytic active state model, revealing a pronounced steric clash between the insertion loop of BIRC6^UBC^ and the Gate helix of SpUba1. **f,** Structural comparison between the pre-transthioesterification and catalytic active states of the BIRC6^UBC^–UBA1 complex reveals that steric clash between the insertion loop and gate helix prevents conformational transition.

Next, we docked the BIRC6^UBC^ (from our resolved UBA6–BIRC6^UBC^–Ub^T^ structure) into the reported doubly loaded UBA1–Ub^T^-Ub^A^ structure (PDB: 4NNJ) by aligning with the UFD domain of UBA1 and UBA6 (**Fig. 5d**) ^58^. This yielded a composite model with BIRC6^UBC^ bound to doubly loaded UBA1–Ub^T^-Ub^A^ in which the UFD domain adopts a distal conformation (hereafter as pre-transthioesterification state). The pre-transthioesterification state indicates that the active site of BIRC6^UBC^ is more than 23 Å away from the active site of UBA1 (**Fig. 5d**). Comparing the pre-transthioesterification state to the resolved catalytic transthioesterification state of UBA6–BIRC6^UBC^– Ub^T^ reveals a notable rotation of the UFD, which brings BIRC6^UBC^ closer to UBA1’s catalytic active site (**Fig. 5e**). However, the gate helix of UBA1 is positioned directly in the path of BIRC6^UBC^’s rotation toward the catalytic cysteine (**Fig. 5f**). This implies that the role of gate helix of UBA1 is to restrict BIRC6^UBC^ from transitioning from the pre-transthioesterification to the catalytic active conformation.

Collectively, our structural and biochemical data establish a gating model for E1-E2 transthioesterification specificity. BIRC6^UBC^ exhibits comparable binding affinity for both UBA1 and UBA6, probably enabling the formation of pre-transthioesterification conformation. However, the gating helix of UBA1 sterically restricts BIRC6^UBC^’s access to UBA1-activated Ub^T^, preventing Ub^T^ transfer to the E2 catalytic cysteine.

## Discussion

Although previous studies have systematically resolved the structural landscape of the UBA1–E2 transthioesterification cycle, the molecular mechanism by which UBA6, the second Ub activating enzyme with non-redundant physiological functions in the ubiquitin system, selectively recognizes its specific E2 partners remains largely unknown. Here, we elucidated the molecular mechanism underlying the UBA6-specific activation of the giant E2–E3 chimera BIRC6. Using an activity-based Ub^Dha^ probe, we captured and visualized the cryo-EM structures of the transthioesterification intermediate complexes doubly loaded UBA6–BIRC6^UBC^–Ub^T^–Ub^A^ and singly loaded UBA6– BIRC6^UBC^–Ub^T^.

Through structural alignment and biochemical validation, we identified an insertion loop in BIRC6^UBC^ that dictates its specificity for UBA6. In this model, our MST binding assay indicates that BIRC6^UBC^ binds with comparable affinity to both UBA1 and UBA6 and can likely be recruited by doubly loaded forms of either E1. However, a steric clash occurs between the insertion loop of BIRC6^UBC^ and the gate helix of UBA1, which likely restricts access of the catalytic cysteine in BIRC6^UBC^ to the thioester bond in UBA1~Ub, thereby preventing the acceptance of activated ubiquitin from UBA1. In UBA6, the gate helix adopts a constitutively rotated and shifted conformation. It is rotated by approximately 30 degrees and displaced by 19 Å relative to its position in UBA1. This structural rearrangement creates a complementary cavity that accommodates the insertion loop of BIRC6^UBC^ without steric clash. The insertion loop fits into this binding site in UBA6 like a key in a lock. By contrast, UBA1 lacks a corresponding pocket and cannot support this accommodation (**Fig. 6**).

**Fig. 6.**
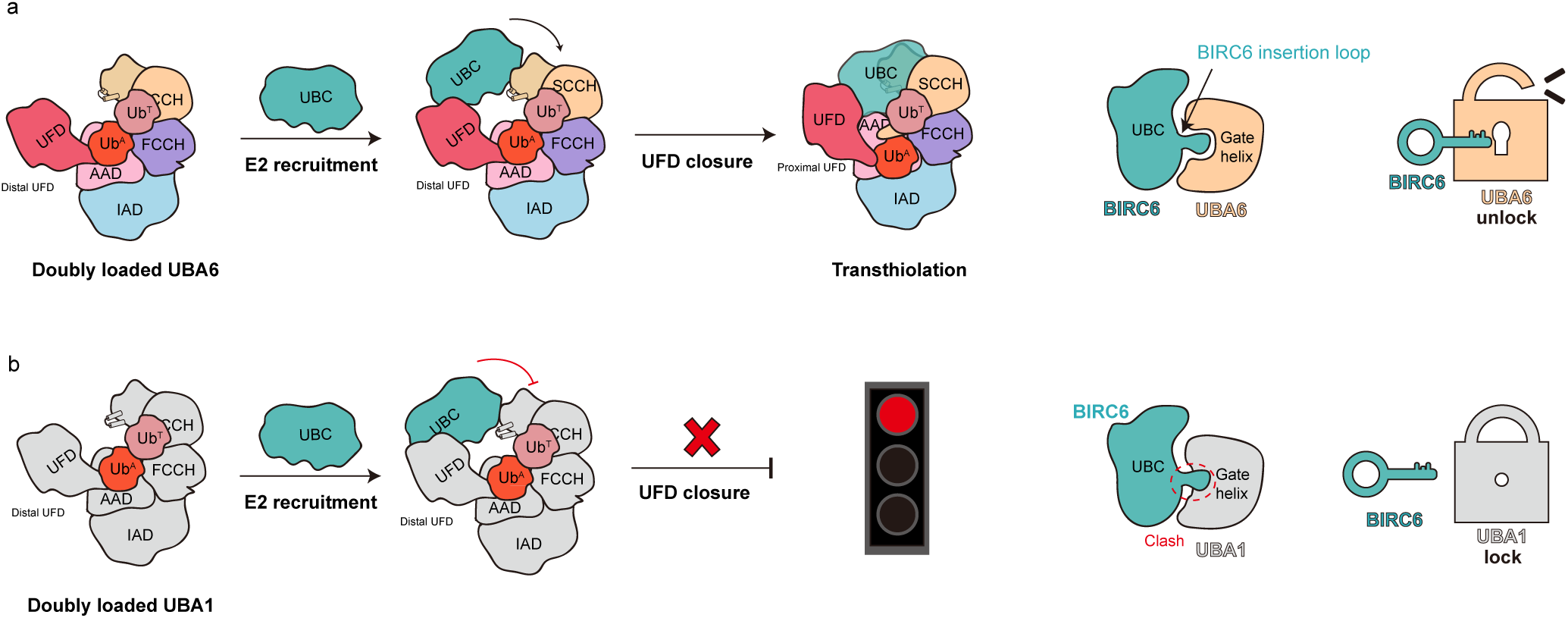
Model of BIRC6^UBC^ specificity with UBA6 over UBA1 in transthioesterification. **a**, UBA6 enables transthioesterification through gate-helix accommodation. In the doubly loaded UBA6-Ub^T^Ub^A^ complex, the UFD domain adopts a closed conformation that positions the catalytic cysteines of UBA6 and BIRC6^UBC^ in proximity for transthioesterification. The insertion loop of BIRC6^UBC^ fits into a compatible cavity created by the rotated gate helix of UBA6, enabling catalysis (unlock). **b**, UBA1 fails to catalyze transthioesterification due to steric clash. Although BIRC6^UBC^ binds UBA1, its insertion loop sterically clashes with the UBA1 gate helix, preventing UFD closure and catalytic alignment. This steric block inhibits transthioesterification, depicted as a locked state (lock).

In addition to BIRC6, UBE2Z represents another E2-E3 chimera enzyme specifically activated by UBA6^10, 24, 59^. Structural and sequence alignments reveal that UBE2Z (PDB:5A4P) also contains a 5-residue insertion loop, which is likely to cause steric clash with the gate helix of UBA1 (**Extended Fig. 9a-d**) ^60^. Therefore, our structural model may also explain the UBA6-specific activation mechanism of UBE2Z. However, the “lock-and-key” model involving the gate helix of E1 and the insertion loop of E2 proposed in our study may not fully explain the molecular mechanism behind the UBA1-specific E2 enzymes, such as UBE2H and UBE2R2. Further structural and biochemical studies will be required to elucidate the precise determinants of E1–E2 specificity in these cases.

Ub^Dha^ is a commercially available or readily synthesized chemical probe that has been used in activity-based profiling to target E1-E2-E3 ubiquitin enzymes. It is suitable for both proteomic analysis and monitoring enzymatic activities in living cells^57^. In this study, we expanded the application of Ub^Dha^ to structural biology by demonstrating its ability to directly and conveniently capture structures of the transthioesterification intermediate within the doubly loaded E1–E2–Ub^T^–Ub^A^ complex. Furthermore, focused 3D classification of Ub^T^ revealed a continuum of conformations depicting ubiquitin transfer from the FCCH domain to the E2 active site, underscoring the strong potential of Ub^Dha^ for resolving dynamic architectures along the ubiquitination cascade. The Ub^Dha^ probe shows strong potential for capturing transthioesterification intermediates of HECT and RBR E3 ligases in complex with E2 and ubiquitin, and holds promise for extension to other ubiquitin-like protein (UBL) systems, highlighting its broad value as a chemical tool for studying UB/UBL cascades.

## Acknowledgements

This study was supported by the National Natural Science Foundation of China (22137005, T2488301, 92253302, 22227810 for L. Liu, and 32501108 for H. Ai), the National Key R&D Program of China (No. 2022YFC3401500, for L. Liu), the New Cornerstone Science Foundation (for L. Liu), the Shanghai Natural Science Foundation (No. 25ZR1402193 for H. Ai), the Shanghai Frontiers Science Center of Drug Target Identification and Delivery (No. ZXWH2170101 for H. Ai), the Shanghai Key Laboratory for Antibody-Drug Conjugates with Innovative Target (No. 24dz2261300 for H. Ai). We acknowledge the Tsinghua University Branch of China National Center for Protein Sciences (Beijing) for cryo-EM screening and data collection in 200 kV Arctica Tecnai microscopy, and the Shuimu BioSciences (Hangzhou) for 300kV cryo-EM data collection.

## Author Contributions

Z. Tong, H. Ai, and L. Liu proposed the idea, designed the experiments and analyzed the results. R. Yuan, Z. Tong, X. Wu cloned the plasmids and expressed the proteins. R. Yuan synthesized the Ub^Dha^ probe. Z. Tong, R. Yuan prepared the cryo-EM samples. Z. Tong collected the cryo-EM data and processed the cryo-EM data, and built the atomic models. R. Yuan, Z. Tong and X. Wu performed the biochemical assay. Z. Tong, R. Yuan, and X. Wu collated the experimental data and prepared the figure panels and tables. Z. Tong. drafted the manuscript. Z. Tong, R. Yuan, X. Wu, H. Ai, and L. Liu revised the manuscript. All authors (Z. Tong, R. Yuan, X. Wu, H. Cai, Z. Xu, H. Ai, and L. Liu) read, discussed and analyzed the manuscript. H. Ai and L. Liu supervised the project.

## Methods

### Plasmids and molecular cloning

The pDARMO-3×FLAG-BIRC6 expression plasmid was purchased from Addgene (plasmid #197967, deposited by Eric Fischer). The cDNA of human UBA6 (Uniprot entry A0AVT1, full-length, amino acid residues 1-1052), BIRC6 UBC domain (Uniprot entry Q9NR09, amino acid residues 4520-4857) were synthesized with sequence optimization for *Escherichia coli* recombinant expression by GenScript Biotech (Nanjing, China). The UBA6 sequence was subsequently cloned into a pDARMO_CMVT vector with an N-terminal 3×FLAG tag. The BIRC6 UBC domain sequence was cloned into the pET-28a vector with an N-terminal His_6_ tag followed by an HRV 3C protease cleavage site. Plasmids of human UBA1 and human ubiquitin were constructed in accordance with previous reports. The generation of protein mutants and truncations was achieved through standard site-directed PCR mutagenesis or homologous recombination techniques and verified by DNA sequencing (Tsingke, Beijing).

### Protein expression and purification

BIRC6 and UBA6 were expressed in HEK293F cells. The cells were cultivated in Union-293 chemically defined medium (Union-Biotech, UP10000) and transfected using polyethyleneimine (PEI, Polysciences) at a density of 1.5-2.0×10^6^ cells·mL^−1^. After 60 h of transfection, the transfected cells were harvested at 5,000 g, resuspended in lysis buffer (50 mM HEPES, pH 7.4, 200 mM NaCl, 5% (v/v) glycerol) supplemented with EDTA-free cOmplete protease inhibitor tablets (Roche, 04693132001). Cells were lysed by sonication and centrifuged at 30,000 g for 45 min at 4 °C. The cell pellets were incubated with anti-DYKDDDDK affinity beads (Smart-Lifesciences, SA042025) for 2 h at 4 °C. The elution of bound proteins was conducted using lysis buffer supplemented with 1 mg·mL^-1^ FLAG peptide (homemade). The eluted 3×FLAG-tagged proteins were then subjected to further purification using a 5 mL prepacked Hitrap Q HP column (Cytiva #17115301). Peak fractions were concentrated and further purified using size-exclusion chromatography (SEC, Superose 6 Increase 10/300 GL or Superdex 200 Increase 10/300 GL, GE Healthcare) in SEC buffer (30 mM HEPES, pH 7.4, 150 mM NaCl).

UBA1 and BIRC6^UBC^ were expressed in *Escherichia coli* BL21 (DE3) cells (Transgene). Cells were cultivated at 37 °C in Luria-Bertani (LB) medium (Coolaber, Cat#PM0010) to an OD_600_ of approximately 0.8 and induced overnight with 0.4 mM isopropyl β-D-thiogalactopyranoside (IPTG) (purchased from LABLEAD Inc., Cat#0487) at 16 °C. After harvesting, the cell pellets were resuspended in lysis buffer (50mM HEPES, pH 7.4, 150 mM NaCl) supplemented with 1 mM phenylmethylsulfonyl fluoride (PMSF). Cells were lysed by sonication and centrifuged at 30,000 g for 30 min at 4 °C, the resulting supernatant was incubated with Ni–NTA affinity resin (purchased from LABLEAD Inc., Cat# N30210) at 4 °C for 1 h. Bound proteins were subsequently eluted using lysis buffer supplemented with 300 mM imidazole. For BIRC6^UBC^, HRV 3C protease was added to remove the His_6_ tag, and the protein was subsequently purified using a Superdex™ 75 Increase 10/300 GL (GE Healthcare) in SEC buffer (50 mM HEPES, pH 7.4, 150 mM NaCl). For UBA1, the protein was purified by ion exchange chromatography using a 5 mL Hitrap Q column. Peak fractions were concentrated and further purified using SEC on a Superdex 200 column (GE Healthcare).

### Fluorescent labeling of ubiquitin

MCQ-Ub (1 equivalent, a ubiquitin variant with an additional cysteine inserted between the N-terminal methionine and glutamine) was concentrated to approximately 2 mM in reaction buffer (20 mM HEPES, pH 7.5, 150 mM NaCl, 0.5mM Tris (2-carboxyethyl) phosphine (TCEP, Aladdin)). Subsequently, 2 equivalents of Oregon Green™ 488 (OG488) maleimide (dissolved in dimethyl sulfoxide (DMSO), purchased from Thermo Fisher Scientific, O6034) were added, and the pH was adjusted to 7.4. The above reaction was conducted at 37 °C in dark for 1 h, and then quenched by the addition of 50 mM dithiothreitol (DTT). The reaction was monitored by analytical reversed-phase high-performance liquid chromatography (RP-HPLC). The final product, fluorescently labeled ubiquitin (Ub^OG488^), was purified using a Superdex 75 Increase column (GE Healthcare) in SEC buffer (50 mM HEPES, pH 7.4, 150 mM NaCl).

### Preparation of Ub^Dha^

UbG76C (1 equivalent, a ubiquitin variant with C-terminal glycine 76 to cysteine mutation) was dissolved in reaction buffer (6 M Gn-HCl, 100 mM NaH_2_PO_4_, pH 9.0), and then DTT was added to a final concentration of 0.5 mM. 50 equivalents of α,α”-di-bromo-adipyl(bis)amide (dissolved in DMSO) was added into the above reaction, and the pH was adjusted to 9.5. The reaction was conducted at 37 °C for 2.5 h. The final product was separated by semi-preparative HPLC (214 nm) and characterized by ESI-MS.

After lyophilizing, Ub^Dha^ was dissolved in buffer A (8 M urea, 20 mM HEPES, pH 7.4, 150 mM NaCl). Subsequently, buffer B (20 mM HEPES, pH 7.4, 150 mM NaCl) was slowly added and thoroughly mixed until the concentration of urea was diluted to 0.5 M. The refolded Ub^Dha^ probe was further purified using a Superdex 75 Increase column (GE Healthcare) pre-equilibrated with SEC buffer (50 mM HEPES, pH 7.4, 150 mM NaCl).

### Reconstitution of UBA6-BIRC6^UBC^-Ub transthioesterification mimic

For the preparation of UBA6-BIRC6^UBC^-Ub transthioesterification mimic, 1 µM UBA6, 1 µM BIRC6^UBC^ and 12.5 µM Ub^Dha^ were mixed in the reaction buffer (50 mM HEPES, pH 7.4, 150 mM NaCl, 5 mM MgCl_2_ and 10 mM ATP) at 37 °C for 1 h. The reaction was monitored through SDSPAGE. The trapped product was further separated using a Superdex 200 Increase 10/300 GL column (GE Healthcare) in SEC buffer (50 mM HEPES, pH 7.4, 150 mM NaCl). Peak fractions were concentrated to 4.7µM for plunging.

### Cryo-EM sample preparation and data collection

3.5 µL prepared complex was added into the glow-discharged holey gold grids (Quantifoil R1.2/1.3, Au mesh) in a gradual manner, and incubated at 4 °C under 100% humidity for 60 seconds. Grids were rapidly immersed into liquid ethane by using the Vitrobot (Thermo Fisher Scientific, blot time 3s, blot force 3).

A total of 21,555 cryo-EM micrographs of the UBA6-BIRC6^UBC^-Ub^Dha^ dataset were collected on a 300 kV Titan Krios G4 microscope configured with a Falcon4 direct electron detector camera using the EPU software (Thermo Fisher Scientific). Micrographs for this dataset were acquired at 75,000× magnification, corresponding to a pixel size of 1.059 Å, under identical exposure conditions. Each image was recorded over 32 frames within 5.59 seconds, with defocus values ranging from −1.0 to −2.0 μm.

### Cryo-EM image processing

All datasets were processed using RELION v3.1.1^61^. The following procedures were performed: motion correction^62^, contrast transfer function (CTF) estimation^63^, manual or automated particle picking, and particle extraction. Subsequent steps in the methodology comprised iterative 2D and 3D classifications, mask generation, 3D auto-refinement, and postprocessing. The data processing flow charts are detailed in **Extended Data Fig. 2**. The statistics pertaining to the processing of cryo-EM data have been summarized in **Table 1**. The determination of overall resolutions was based on the gold standard Fourier shell correlation (FSC) criterion at 0.143, and local resolution estimations were carried out using ResMap v1.1.4.

**Table 1.** Cryo-EM data collection, refinement, and validation statistics.

| **Table 1. Cryo-EM data collection, refinement, and validation statistics** |  |  |
| --- | --- | --- |
|  | Singly loaded UBA6–BIRC6<sup>UBC</sup>–Ub<sup>T</sup> complex (EMD- 66842, PDB 9XGB) | Doubly loaded UBA6–BIRC6<sup>UBC</sup>–Ub<sup>T</sup>–Ub<sup>A</sup> complex (EMD- 66843, PDB 9XGC) |
| Magnification | 96000 | 96000 |
| Voltage (kV) | 300 | 300 |
| Spherical aberration (mm) | 2.7 | 2.7 |
| Detector | K3 | K3 |
| Electron exposure (e<sup>-</sup>/Å<sup>2</sup>) | 51.35 e<sup>-</sup>, 32 frames | 51.35 e<sup>-</sup>, 32 frames |
| Defocus range (μm) | -1.5 to -2.0 | -1.5 to -2.0 |
| Pixel size (Å) | 0.808 | 0.808 |
| Symmetry imposed | C1 | C1 |
| Box size (pixel) | 200 | 200 |
| Micrographs (no.) | 21555 | 21555 |
| Initial particle images (no.) | 14,438,711 | 14,438,711 |
| Final particle images (no.) | 1,201,224 | 1,087,360 |
| Map resolution (Å) | 3.3 Å | 3.4 Å |
| Map resolution range | 2.9–5.8 Å | 2.9–5.8 Å |
| FSC threshold | 0.143 | 0.143 |
| Initial model use (PDB code) | 1UBQ, 3CEG, 7PVN | 1UBQ, 3CEG, 7PVN |
| Model resolution (Å) | 4.0 Å | 3.5 Å |
| FSC threshold | 0.5 | 0.5 |
| Map sharpening B factor (Å<sup>2</sup>) | -180 | -180 |
| Nonhydrogen atoms | 10826 | 11421 |
| Protein residues | 1366 | 1440 |
| Nucleotides | 0 | 0 |
| Protein | 30.00/534.77/209.50 | 131.16/461.09/237.04 |
| Nucleotide | --- |  |
| Bond lengths (Å) | 0.009 | 0.008 |
| Bond angles (°) | 1.428 | 1.142 |
| MolProbity score | 1.65 | 1.71 |
| Clashscore | 6.58 | 7.8 |
| Poor rotamers (%) | 1 | 0.08 |
| Favored (%) | 95.8 | 95.86 |
| Allowed (%) | 4.12 | 4.14 |
| Disallowed (%) | 0.07 | 0 |

### Model building and refinement

The cryo-EM density maps of the singly loaded UBA6–BIRC^6UBC^–Ub^T^ complex and the doubly loaded UBA6–BIRC6^UBC^–Ub^T^–Ub^A^ complex were employed for structural modeling. In the case of the singly loaded UBA6–BIRC6^UBC^–Ub^T^ assembly, the atomic coordinates of BIRC6^UBC^ (PDB: 3CEG)^59^, ubiquitin (PDB: 1UBQ)^64^, and UBA6 (PDB: 7PVN) ^11^ were fitted as rigid bodies into the cryo-EM density using ChimeraX-1.7.13949^65^. The resulting coordinates, which preserved spatial positioning, were subsequently imported into WinCoot-0.8.240, where BIRC6^UBC^, Ub^T^, and UBA6 were combined into a single initial model^66^.

For the doubly loaded UBA6–BIRC6UBC–Ub^T^–Ub^A^ complex, the structure of the singly loaded UBA6–BIRC6UBC–Ub^T^ was manually positioned together with Ub^A^ in ChimeraX-1.7.1, followed by integration of Ub^A^ in WinCoot-0.8.240 to produce the starting model. Initial models were refined in Phenix-1.19.254 through real-space refinement, incorporating secondary structure constraints and geometric restraints^67^. All models underwent manual inspection and adjustment in COOT-0.8.2.55. A summary of the 3D reconstruction and modeling statistics is provided in **Table 1**. Structural visualizations were generated in ChimeraX-1.7.139, from which figures were exported for presentation.

### E1~Ub thioester formation assays

E1~Ub thioester formation assays were performed with 25 µM Ub^OG488^, 0.2 µM E1 (UBA1, UBA6, or E1 mutants) in Ubiquitylation Buffer (50 mM HEPES, pH 7.4, 150 mM NaCl, 5 mM MgCl_2_ and 10 mM ATP). Reactions were incubated at 37 °C for 30 min and quenched with non-reducing protein loading buffer, followed by analysis on 4–12% SurePAGE™ Bis-Tris gels (GenScript Biotech) and fluorescence visualization.

### E1-E2 thioester transfer assays

E1-E2 thioester transfer assays were performed with 25 µM Ub^OG488^, 0.2 µM E1 (UBA1, UBA6, or E1 mutants) and either 0.2 µM BIRC6 or 5 µM BIRC6^UBC^ (or its mutants) in Ubiquitylation Buffer (50 mM HEPES, pH 7.4, 150 mM NaCl, 5 mM MgCl_2_ and 10 mM ATP). To assess critical structural interfaces, reactions were incubated at RT for 2 min and quenched with non-reducing protein loading buffer. To validate the molecular basis of Uba6 specificity for BIRC6^UBC^, reactions were incubated at 37 °C for 30 min and quenched with non-reducing protein loading buffer. Samples were resolved on 3–8% NuPAGE Bis-Tris gels (Thermo Fisher Scientific) or 4–12% SurePAGE™ Bis-Tris gels (GenScript Biotech), and then imaged by fluorescence or Coomassie Brilliant Blue staining.

### E1 competitive assays

For E1 competitive assays, 40 µM Ub^OG488^, 0.1 µM UBA6 and 5 µM BIRC6^UBC^ were first mixed in the reaction buffer (50 mM HEPES, pH 7.4, 150 mM NaCl, 2 mM MgCl_2_). The UBA1 (final concentrations of 0/1/2/4/6 µM) was then added to the mixture, followed by the addition of 4× ATP buffer (25 mM MgCl_2_, pH 7.5, 50 mM ATP) to initiate the reactions. Reactions were incubated at 37 °C for 2 min and quenched with non-reducing protein loading buffer. Samples were resolved on 4-12% SurePAGE™ Bis-Tris gels (GenScript Biotech), and then imaged by fluorescence or Coomassie Brilliant Blue staining.

### MicroScale Thermophoresis (MST) analysis

UBA1 and UBA6 were labelled with the His-tag labelling Kit RED-tris-NTA 2^nd^ Generation dye (Nano temper Cat# MO-L018). Based on preliminary measurements, the affinity of UBA1 towards the dye was determined to be greater than 10 nM. Then, 100 nM dye and 200 nM UBA1 were used as per manufacturer’s instructions. Conversely, the affinity of UBA6 towards the dye was determined to be weaker than 10 nM. Therefore, 100 nM dye and 800 nM UBA6 were used as per manufacturer’s instructions. For the binding assay, a serial dilution of the diluted BIRC6^UBC^ (from 1.2 mM to 0.01724 µM) was prepared using the MST buffer (50 mM HEPES, pH 7.4, 150 mM NaCl, 0.05% Tween-20). Then, labelled E1 (100 nM UBA1, or 400 nM UBA6) was added to each group. The affinity measurements were performed on a Monolith NT.115 (NanoTemper) at Medium MST power, with each independent experiment comprising two technical replicates. The binding affinities (Kd) were calculated by MO. Affinity Analysis software (Nano Temper).

## Data availability

Cryo-EM maps have been deposited in the Electron Microscopy Data Bank (EMDB, www.ebi.ac.uk/pdbe/emdb/) under the accession codes EMD-66842 (singly loaded UBA6– BIRC6^UBC^–Ub^T^ complex), EMD-66843 (doubly loaded UBA6–BIRC6^UBC^–Ub^T^-Ub^A^ complex). The atomic models have been deposited in the Protein Data Bank (PDB, www.rcsb.org) under the accession codes 9XGB (singly loaded UBA6–BIRC6^UBC^–Ub^T^ complex) and 9XGC (doubly loaded UBA6–BIRC6^UBC^–Ub^T^-Ub^A^ complex).

## Extended Data Figure Legends

**Extended Data Fig. 1 | Semi-synthesis of Ub^Dha^ probe and preparation of UBA6-BIRC6^UBC^-Ub^Dha^ complex. a**, Schematic of the semi-sythesis of Ub^Dha^ from recombinant UbG76C. **b**, SDS-PAGE analysis of purified Ub^Dha^. **c**, Representative size-exclusion chromatography (SEC) of Ub^Dha^ purification. **d**, SDS-PAGE analysis of the preparation of UBA6-BIRC6^UBC^-Ub^Dha^ complex at indicated times. **e**, Representative size-exclusion chromatography of UBA6-BIRC6^UBC^-Ub^Dha^ complex. **f**, SDS-PAGE analysis of UBA6-BIRC6^UBC^-Ub^Dha^ complex.

**Extended Data Fig. 2 | Cryo-EM data collection and processing of UBA6-BIRC6^UBC^-Ub^Dha^ complexes. a**, A representative micrograph from the UBA6-BIRC6^UBC^-Ub^Dha^ dataset. **b**, CTF estimation of the micrograph shown in **a**. **c**, Representative 2D averages of the UBA6-BIRC6^UBC^Ub^Dha^ complex. **d**, Cryo-EM processing flowchart of the UBA6-BIRC6^UBC^-Ub^Dha^ dataset. The distribution of the Euler angles within UBA6-BIRC6^UBC^-Ub^T^-Ub^A^ complex and UBA6-BIRC6^UBC^Ub^T^ complex have been shown.

**Extended Data Fig. 3 | Cryo-EM density of the UBA6-BIRC6^UBC^-Ub^Dha^ complexes and focused Ub^T^ in UBA6-BIRC6^UBC^-Ub^Dha^ complexes. a**, Fourier shell correlation (FSC) curves of the masked map after Relion postprocessing. The resolution was determined by the FSC=0.143 criterion. **b**, The local resolution of the density of the map calculated by Relion. **c**, Cryo-EM reconstructions and corresponding models of Ub^T^ states resulting from 3D classification without image realignment. A focus mask was applied to the Ub^T^ region in both doubly- and single-loaded complexes. The corresponding orientation of Ub^T^ was shown at right.

**Extended Data Fig. 4 | Cryo-EM density and model of the singly loaded UBA6–BIRC6^UBC^–Ub^T^ complex. a-h.** Representative regions of cryo-EM densities for BIRC6^UBC^ (4520-4858) (**a**), Ub^T^ (176) (**b**), UBA6^SCCH^ (613-888) (**c**), UBA6^UFD^ (943-1051) (**d**), overall structure of singly loaded UBA6– BIRC6^UBC^–Ub^T^ complex (**e**), UBA6^FCCH^ (203-297) (**f**), UBA6^AAD^ (430-612, 889-942) (**g**), UBA6^IAD^ (40-202, 298-429) (**h**).

**Extended Data Fig. 4 | Cryo-EM density and model of the doubly loaded UBA6–BIRC6^UBC^–Ub^T^ complex. a-h.** Representative regions of cryo-EM densities for BIRC6^UBC^ (4520-4858) (**a**), Ub^T^ (176) (**b**), UBA6^SCCH^ (613-888) (**c**), UBA6^UFD^ (943-1051) (**d**), overall structure of singly loaded UBA6– BIRC6^UBC^–Ub^T^ complex (**e**), Ub^T^ (1-76) (**f**), UBA6^FCCH^ (203-297) (**g**), UBA6^AAD^ (430-612, 889-942) (**h**), UBA6^IAD^ (40-202, 298-429) (**i**).

**Extended Data Fig. 6 | Comparison of the canonical UBC domain and BIRC6’s UBC domain. ac**, Cartoon depictions of the canonical UBC scaffold from Ubc4 (**a**, PDB: 3L1Y) and the BIRC6’s UBC domain (**c**, PDB: 3CEG). **b**, Structural alignment of the crystal structure of Ubc4 and BIRC6^UBC^ determined in this study. **d**, Sequence alignment of representative UBC domains, with unique secondary structures from BIRC6^UBC^ highlighted.

**Extended Data Fig. 7 | Analysis of the interactions between UBA6 and BIRC6^UBC^. a**, Close-up view of the interface between BIRC6^UBC^ N-extension (residue D4538) and UBA6^UFD^ (residue P999). **b**, Close-up view of the interface between BIRC6^UBC^ N-extension (residues Q4634, Q4567, A4569) and UBA6^AAD^ (Y576, P547, E572). **c**, Close-up views of the interface between BIRC6^UBC^ β1-β2 loop (residues E4604, R4605, L4606) and UBA6^UFD^ (residues V990, K991, M992). **d**, Close-up view of the interface between the BIRC6^UBC^ N-helix (residues R4576, Q4580, V4583) and UBA6^UFD^ (residue V997). **e**, Close-up views of the interface between the BIRC6^UBC^ α3-insertion (residues P4709, Y4711, R4745) and UBA6^SCCH^ (residues F722, F734). **f**, Close-up views of the interface of the BIRC6^UBC^ Cextension (residues R4745, R4799, T4789) and UBA6^SCCH^ (residues S678, S818, N799). Cryo-EM density and the corresponding structural model are shown, with interacting residues represented as sticks.

**Extended Data Fig. 8 | Structural analysis and validation of the molecular basis of Uba6 specificity for BIRC6^UBC^. a**, Structural alignment of the UBA6-BIRC6^UBC^-Ub complexes (this study) with previously determined UBA1-E2-Ub structures. **b-c**, In vitro E1~Ub thioester formation assays using fluorescent labeled Ub. Reactions were performed with UBA1 or its UBA1^GH*^ mutant (**b**), Uba6 or its Uba6^GH*^ mutant (**c**). Gel images are representative (n=3). **d-g**, Structural alignments reveal steric clashes between the insertion loop of BIRC6^UBC^ and the α22-α23 helix of UBA1 across different UBA1–E2 complexes (PDB codes indicated).

**Extended Data Fig. 9 | Structural and sequence alignments of UBE2Z and BIRC6^UBC^. a,** Sequence alignment of UBA6-specific and UBA1-specific E2s. **b-c**, Structural alignment of the crystal structure of UBE2Z (PDB: 5A4P), Ubc4 (PDB: 4II2) and BIRC6^UBC^ determined in this study. **d**, Steric clash between UBA1 and UBE2Z may involve in the insertion loop of UBE2Z and the α22-α23 helix of UBA1.

