## Extended Data Figures for "Steric Gating Dictates Selective Activation of BIRC6 by UBA6 Over UBA1"

### Extended Fig. 1

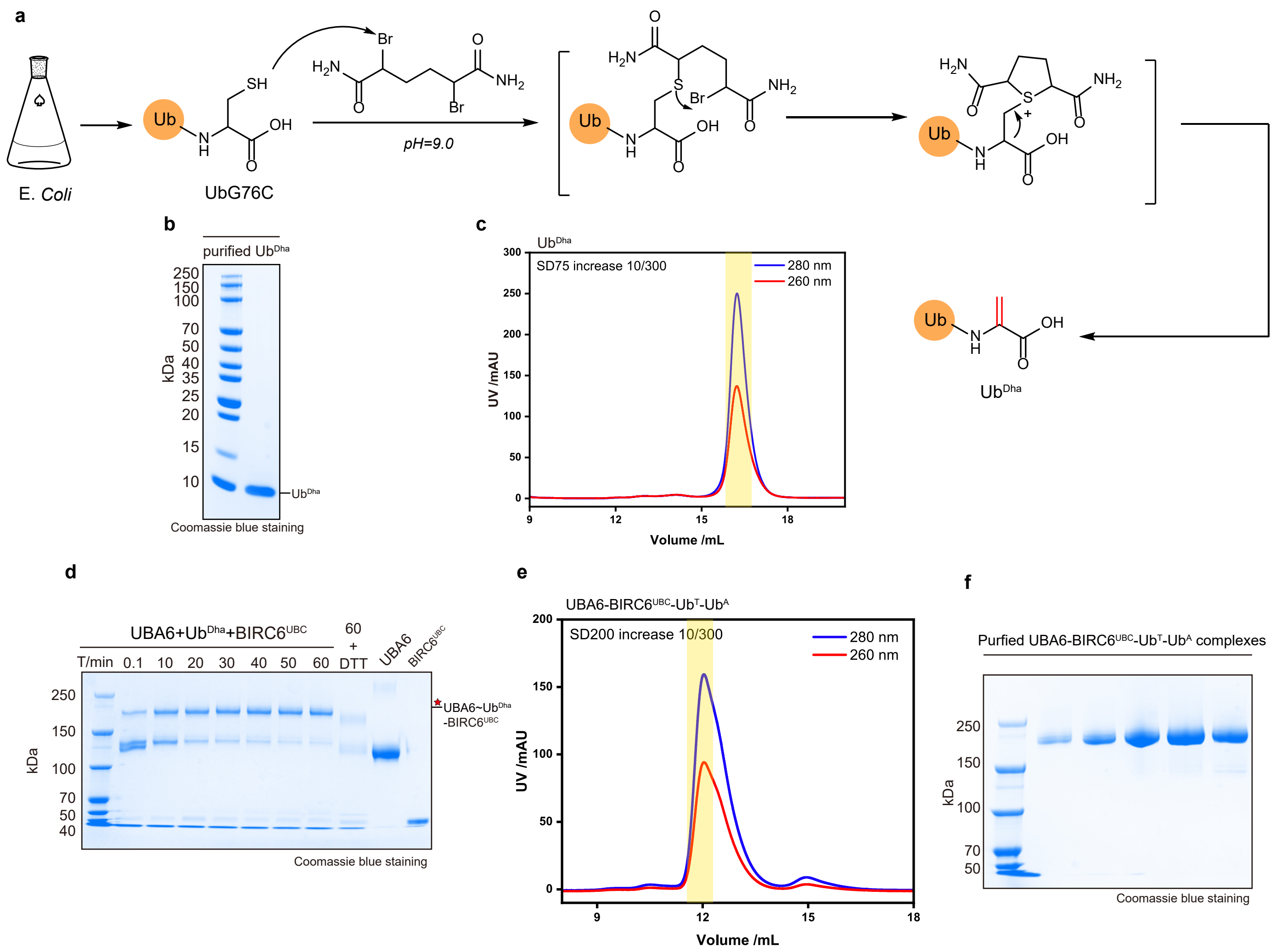


**Extended Data Fig. 1 |** Semi-synthesis of Ub$^{Dha}$ probe and preparation of UBA6-BIRC6$^{UBC}$-Ub$^{Dha}$ complex. **a,** Schematic of the semi-sythesis of Ub$^{Dha}$ from recombinant UbG76C. **b,** SDS-PAGE analysis of purified Ub$^{Dha}$. **c,** Representative size-exclusion chromatography (SEC) of Ub$^{Dha}$ purification. **d,** SDS-PAGE analysis of the preparation of UBA6-BIRC6$^{UBC}$-Ub$^{Dha}$ complex at indicated times. **e,** Representative size-exclusion chromatography of UBA6-BIRC6$^{UBC}$-Ub$^{Dha}$ complex. **f,** SDS-PAGE analysis of UBA6-BIRC6$^{UBC}$-Ub$^{Dha}$ complex.

#### Extended Fig. 2

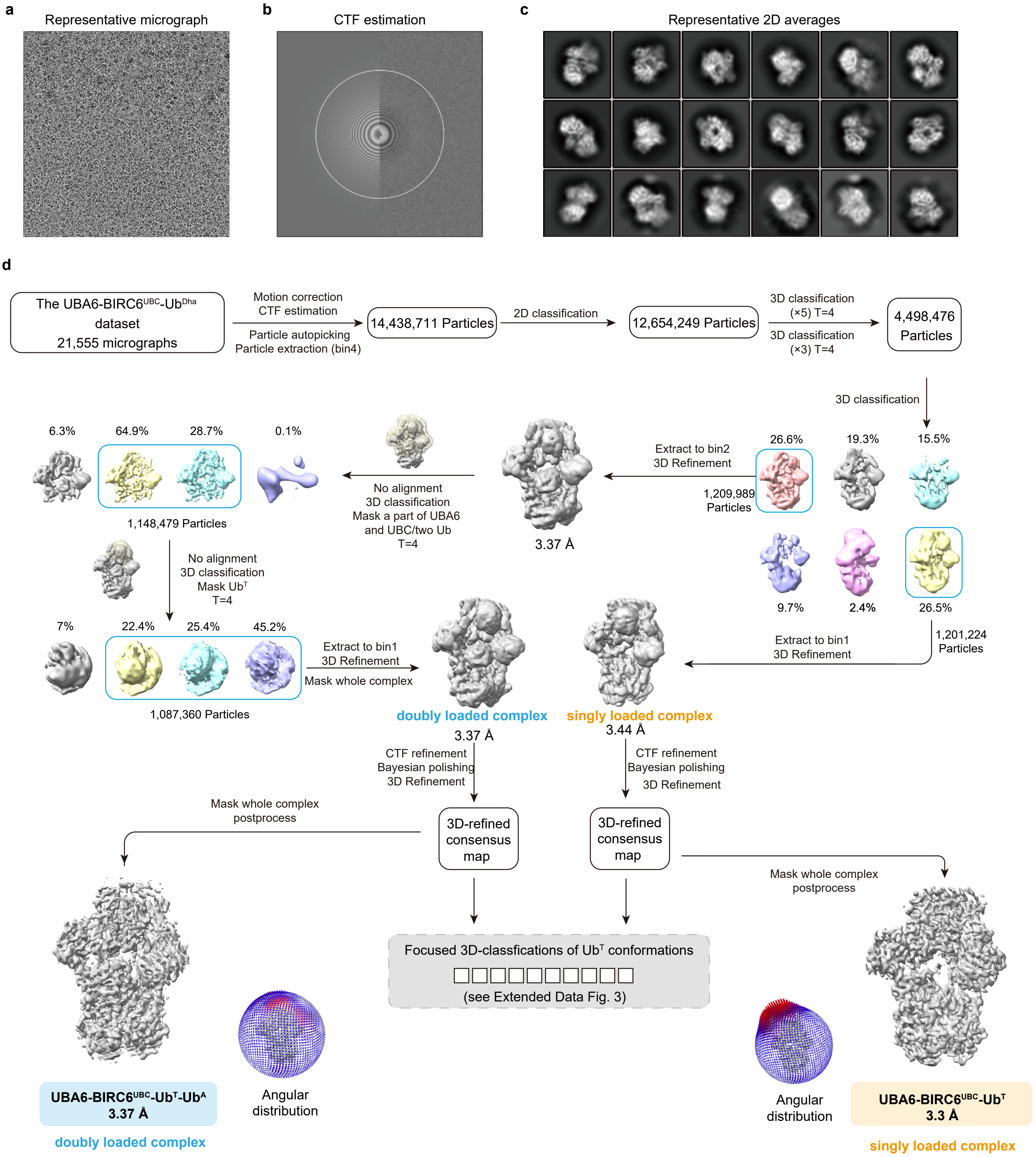


**Extended Data Fig. 2 |** Cryo-EM data collection and processing of UBA6-BIRC6$^{UBC}$-Ub$^{Dha}$ complexes. **a,** A representative micrograph from the UBA6-BIRC6$^{UBC}$-Ub$^{Dha}$ dataset. **b,** CTF estimation of the micrograph shown in **(a)**. **c,** Representative 2D averages of the UBA6-BIRC6$^{UBC}$-Ub$^{Dha}$ complex. **d,** Cryo-EM processing flowchart of the UBA6-BIRC6$^{UBC}$-Ub$^{Dha}$ dataset. The distribution of the Euler angles within UBA6-BIRC6$^{UBC}$-Ub$^{T}$-Ub$^{A}$ complex and UBA6-BIRC6$^{UBC}$-Ub$^{T}$ complex have been shown.

### Extended Fig. 3

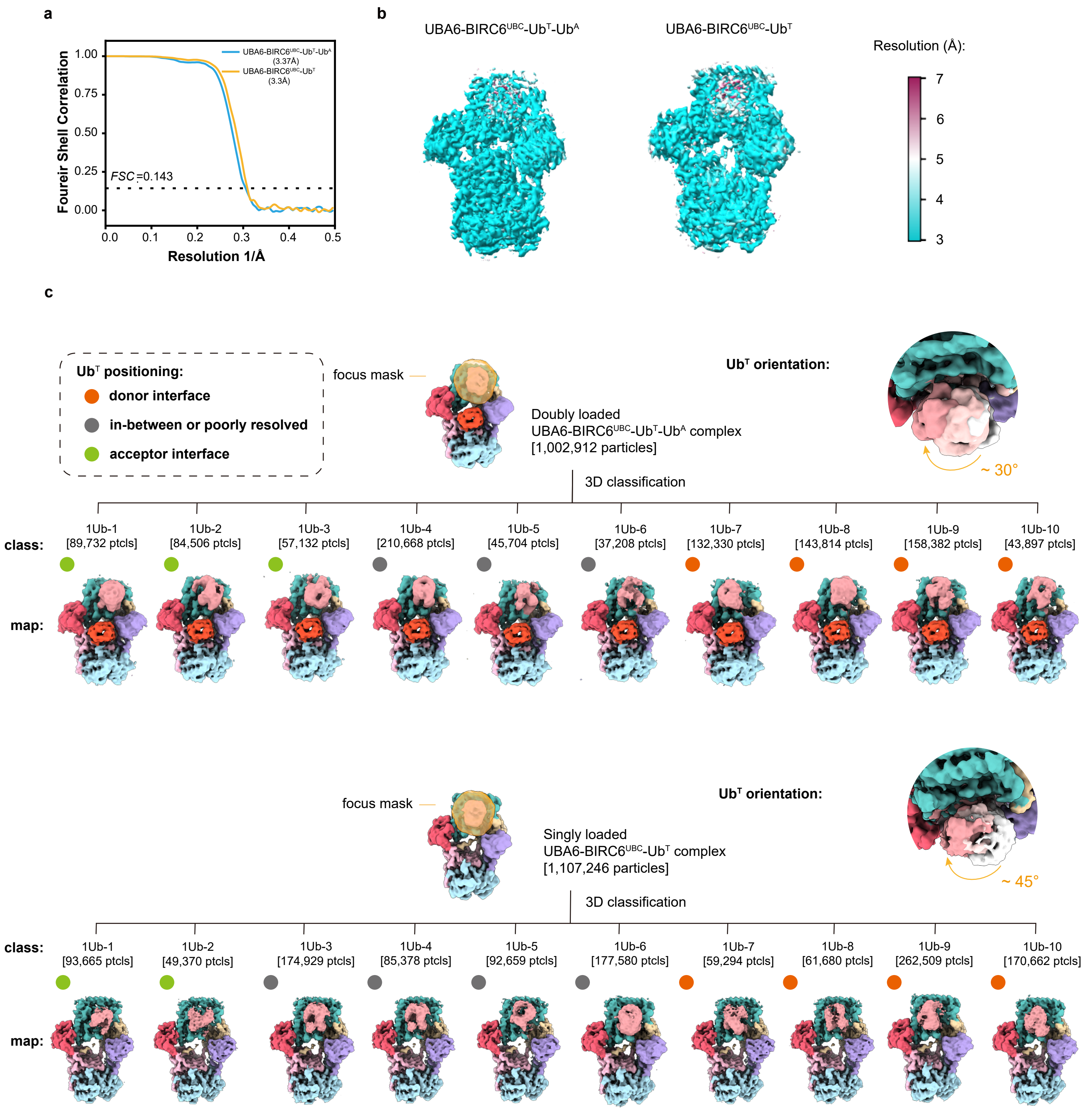


**Extended Data Fig. 3 |** Cryo-EM density of the UBA6-BIRC6$^{UBC}$-Ub$^{Dha}$ complexes and focused Ub$^{T}$ in UBA6-BIRC6$^{UBC}$-Ub$^{Dha}$ complexes. **a,** Fourier shell correlation (FSC) curves of the masked map after Relion postprocessing. The resolution was determined by the FSC=0.143 criterion. **b,** The local resolution of the density of the map caculated by Relion. **c,** Cryo-EM reconstructions and corresponding models of Ub$^{T}$ states resulting from 3D classification without image realignment. A focus mask was applied to the Ub$^{T}$ region in both doubly- and singly-loaded complexes. The corresponding orientation of Ub$^{T}$ were shown at right.

### Extended Fig. 4

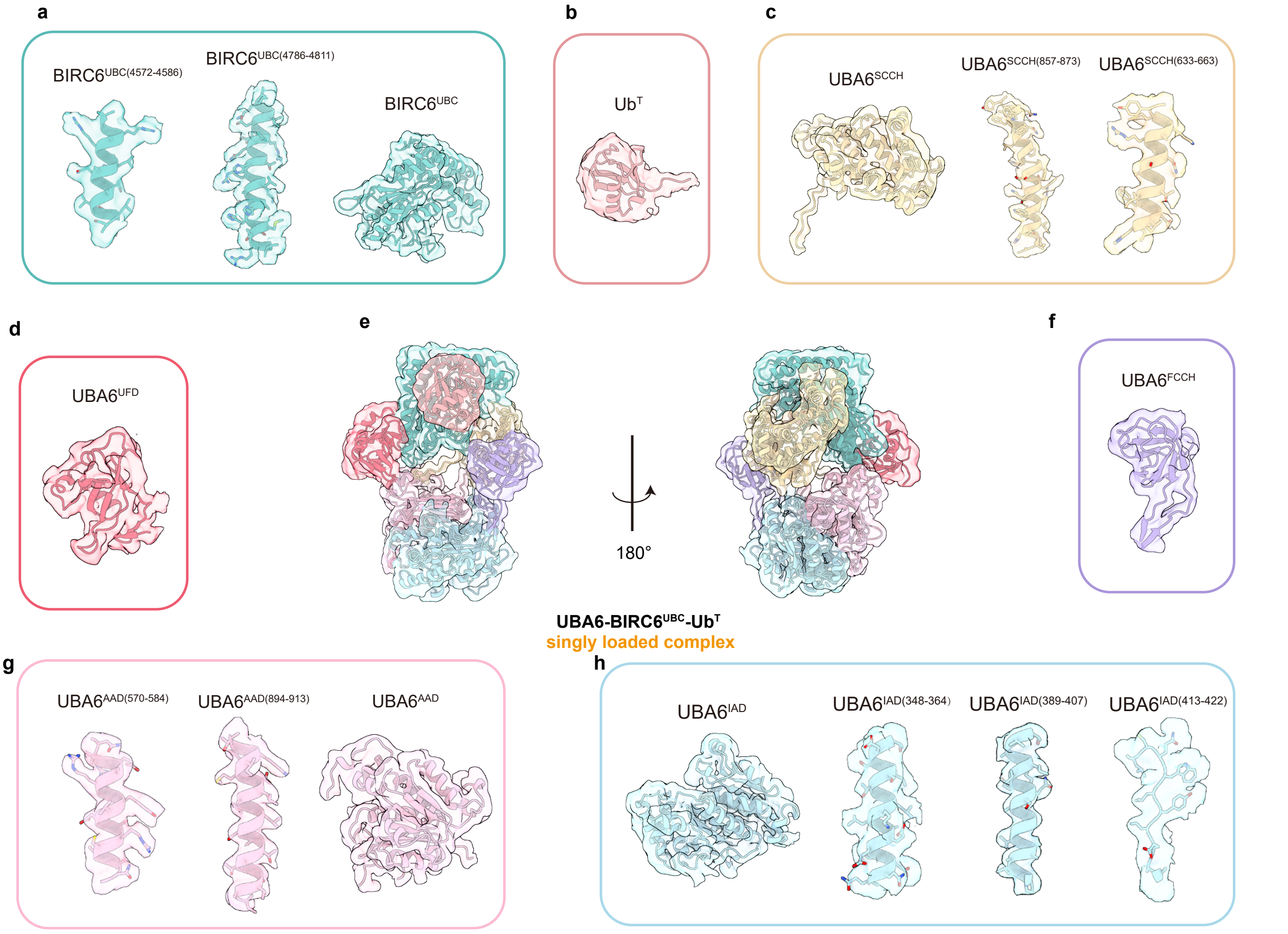


**Extended Data Fig. 4 |** Cryo-EM density and model of the singly loaded UBA6–BIRC6$^{UBC}$–Ub$^{T}$ complex. **a-h.** Representative regions of cryo-EM densities for BIRC6$^{UBC}$ (4520-4857) **(a)**, Ub$^{T}$ (1-76) **(b)**, UBA6$^{SCCH}$ (613-888) **(c)**, UBA6$^{UFD}$ (943-1052) **(d)**, overall structure of singly loaded UBA6–BIRC6$^{UBC}$–Ub$^{T}$ complex **(e)**, UBA6$^{FCCH}$ (203-297) **(f)**, UBA6$^{AAD}$ (430-612, 889-942) **(g)**, UBA6$^{IAD}$ (40-202, 298-429) **(h)**.

#### Extended Fig. 5

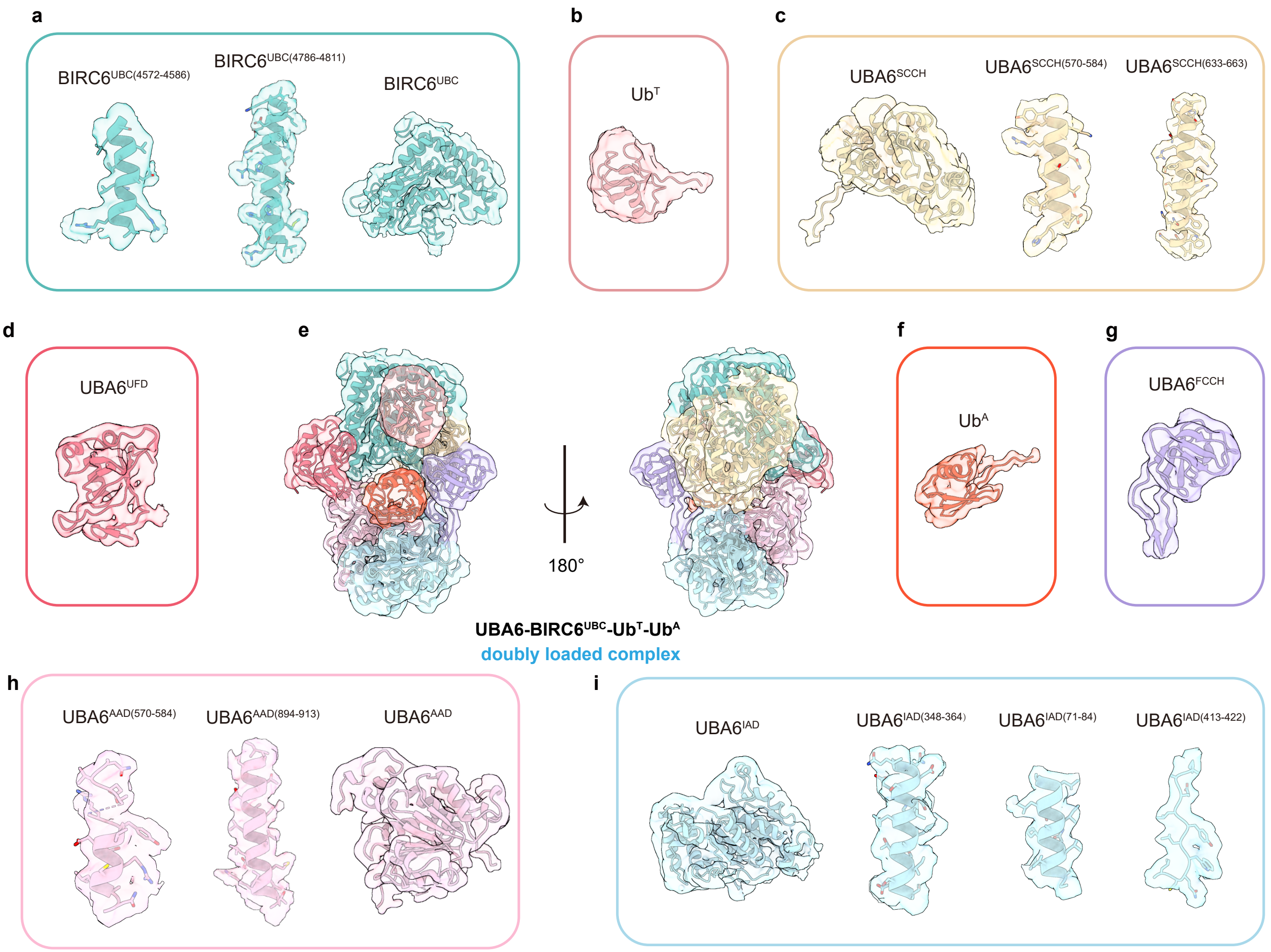


**Extended Data Fig. 5 |** Cryo-EM density and model of the doubly loaded UBA6–BIRC6$^{UBC}$–Ub$^{T}$-Ub$^{A}$ complex. **a-h.** Representative regions of cryo-EM densities for BIRC6$^{UBC}$ (4520-4857) **(a)**, Ub$^{T}$ (1-76) **(b)**, UBA6$^{SCCH}$ (613-888) **(c)**, UBA6$^{UFD}$ (943-1052) **(d)**, overall structure of doubly loaded UBA6–BIRC6$^{UBC}$–Ub$^{T}$-Ub$^{A}$ complex **(e)**, Ub$^{A}$ (1-76) **(f)**, UBA6$^{FCCH}$ (203-297) **(g)**, UBA6$^{AAD}$ (430-612, 889-942) **(h)**, UBA6$^{IAD}$ (40-202, 298-429) **(i)**.

### Extended Fig. 6

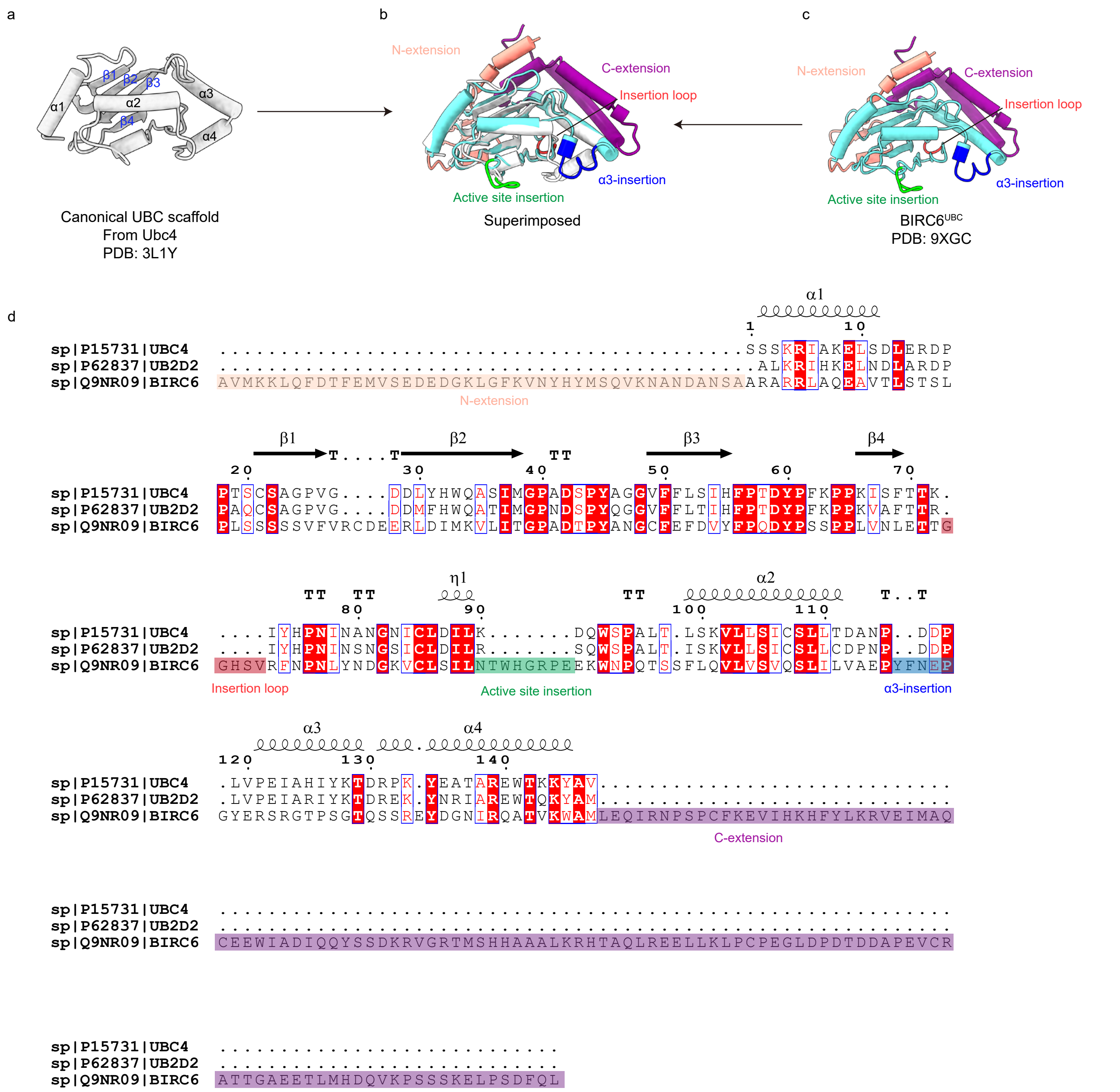


**Extended Data Fig. 6 |** Comparison of the canonical UBC domain and BIRC6's UBC domain. **a-c,** Cartoon depictions of the canonical UBC scaffold from Ubc4 (**a**, Protein Data Bank (PDB): 3L1Y) and the BIRC6's UBC domain (**c**, PDB: 9XGC). **b,** Structural alignment of the crystal structure of Ubc4 and BIRC6$^{UBC}$ determined in this study. **d,** Sequence alignment of representative UBC domains, with unique secondary structures from BIRC6$^{UBC}$ highlighted.

### Extended Fig.7

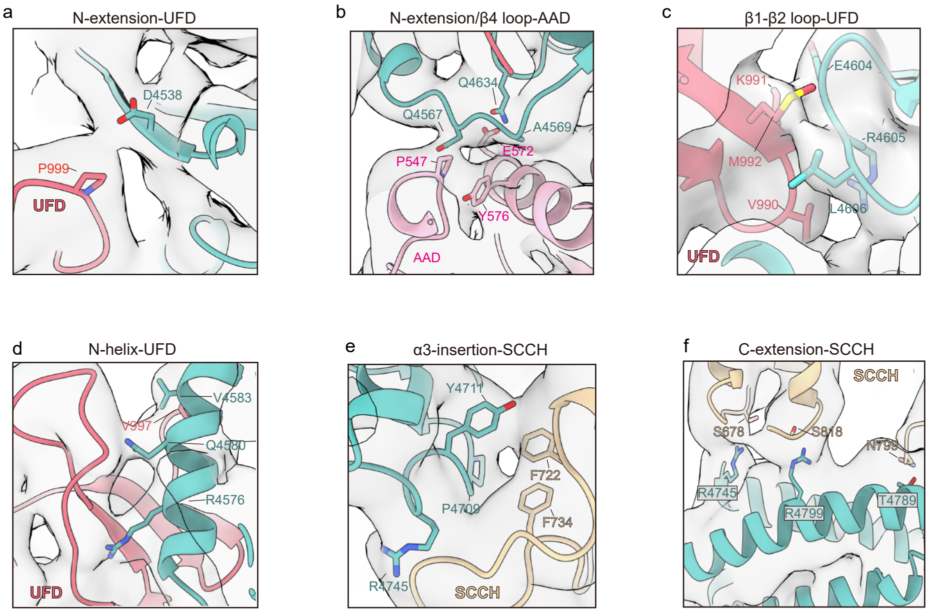


**Extended Data Fig. 7 |** Analysis of the interactions between UBA6 and BIRC6$^{UBC}$. **a,** Close-up view of the interface between BIRC6$^{UBC}$ N-extension (residue D4538) and UBA6$^{UFD}$ (residue P999). **b,** Close-up view of the interface between BIRC6$^{UBC}$ N-extension (residues Q4634, Q4567, A4569) and UBA6$^{AAD}$ (residues Y576, P547, E572). **c,** Close-up views of the interface between BIRC6$^{UBC}$ β1-β2 loop (residues E4604, R4605, L4606) and UBA6$^{UFD}$ (residues V990, K991, M992). **d,** Close-up view of the interface between the BIRC6$^{UBC}$ N-helix (residues R4576, Q4580, V4583) and UBA6$^{UFD}$ (residue V997). **e,** Close-up views of the interface between the BIRC6$^{UBC}$ α3-insertion (residues P4709, Y4711, R4745) and UBA6$^{SCCH}$ (residues F722, F734). **f,** Close-up views of the interface of the BIRC6$^{UBC}$ C-extension (residues R4745, R4799, T4789) and UBA6$^{SCCH}$ (residues S678, S818, N799). Cryo-EM density and the corresponding structural model are shown, with interacting residues represented as sticks.

### Extended Fig. 8

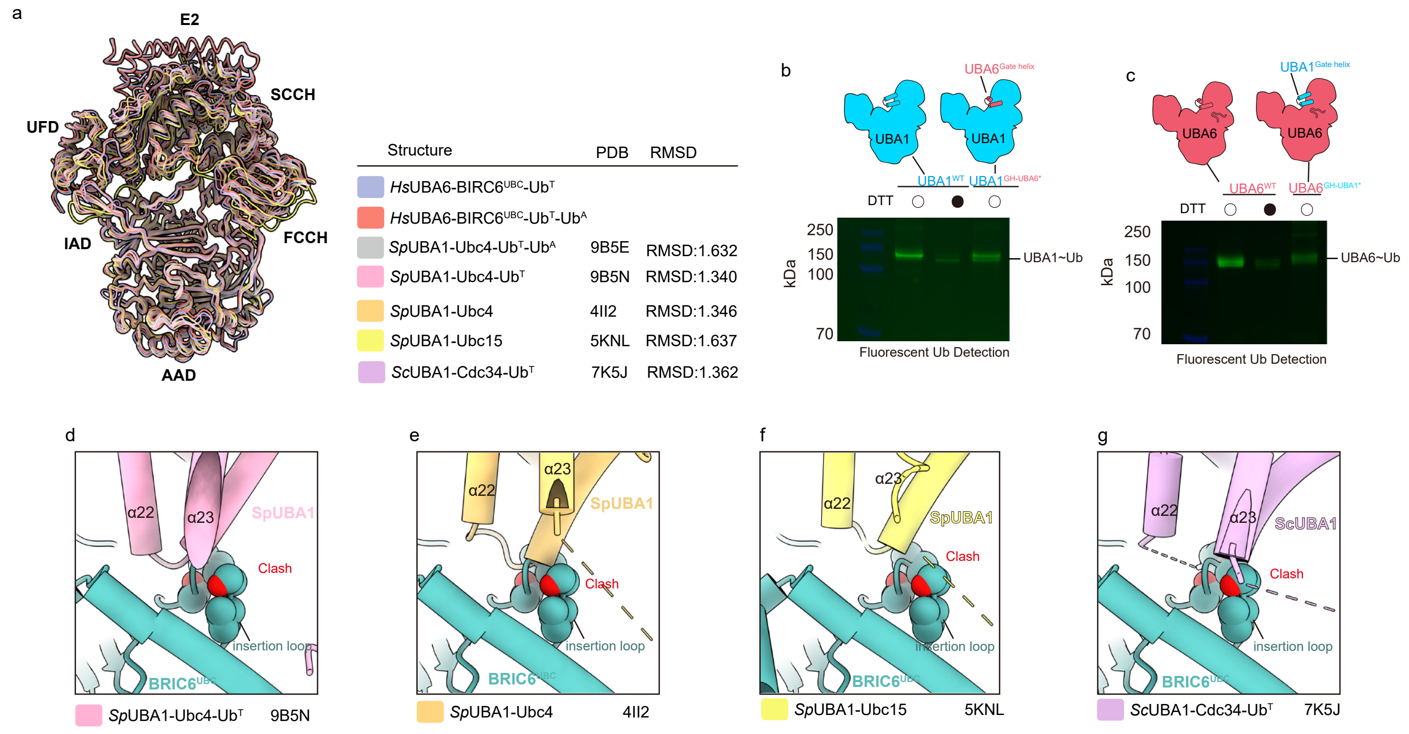


**Extended Data Fig. 8 |** Structural analysis and validation of the molecular basis of UBA6 specificity for BIRC6$^{UBC}$. **a,** Structural alignment of the UBA6-BIRC6$^{UBC}$-Ub complexes (this study) with previously determined UBA1-E2-Ub structures. **b-c,** In vitro E1~Ub thioester formation assays using fluorescent labeled Ub. Reactions were performed with UBA1 or its UBA1$^{GH\text{-}UBA6^*}$ mutant **(b)**, UBA6 or its UBA6$^{GH\text{-}UBA1^*}$ mutant **(c)**. Gel images are representative (n=3). **d-g,** Structural alignments reveal steric clashes between the insertion loop of BIRC6$^{UBC}$ and the α22-α23 helix of UBA1 across different UBA1–E2 complexes (PDB codes indicated).

### Extended Fig. 9

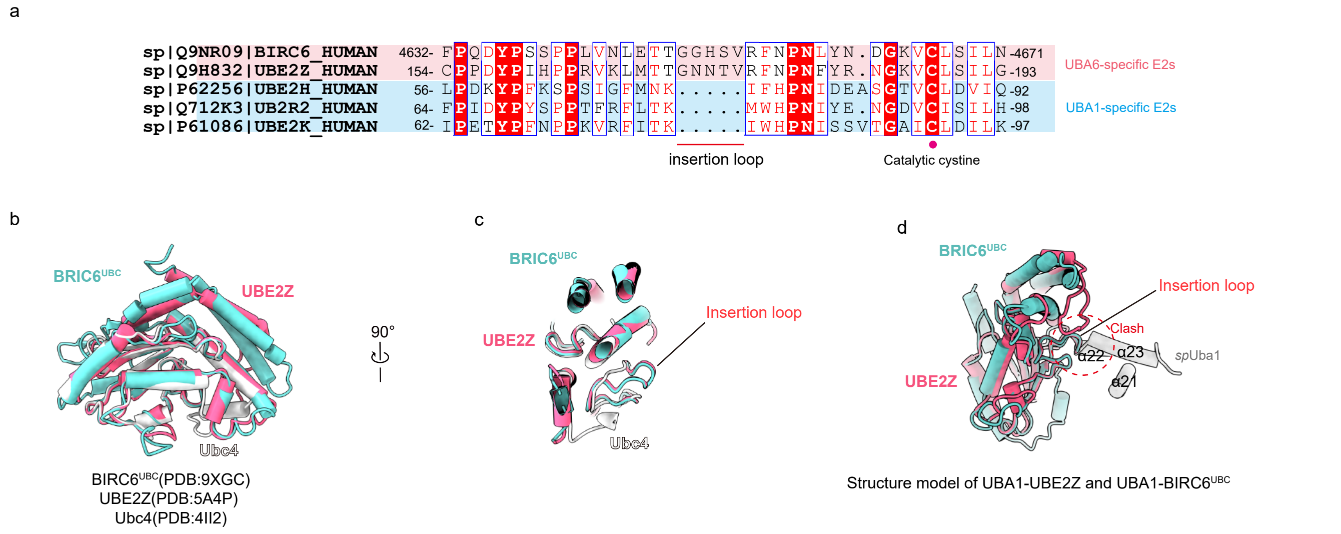


**Extended Data Fig. 9 |** Structural and sequence alignments of UBE2Z and BIRC6$^{UBC}$. **a,** Sequence alignment of UBA6-specific and UBA1-specific E2s. **b-c,** Structural alignment of the crystal structure of UBE2Z (PDB: 5A4P), Ubc4 (PDB: 4II2) and BIRC6$^{UBC}$ determined in this study. **d,** Steric clash between UBA1 and UBE2Z may involve in the insertion loop of UBE2Z and the α22-α23 helix of UBA1.
